# Leveraging whole genome sequencing to promote genetic diversity and population health in zoo-housed western lowland gorillas

**DOI:** 10.1101/2023.11.13.565567

**Authors:** J. E. Gorzynski, M. D. Danforth, V. Strong, I. Kutinsky, H. Murphy, L. Lowenstine, J. R. Priest, E. A. Ashley

**Author notes:** Correspondence Euan Ashley MD PhD 870 Quarry Road, Stanford, 94304.

## Abstract

The sustainability of zoo populations is dependent on maintaining genetic diversity and controlling heritable disease. Here, we explore the integration of whole genome sequencing data in the management of the international zoological population of western lowland gorillas, focusing on genetic diversity and heritable diseases. By comparing kinship values derived from classical pedigree mapping and whole genome sequencing, we demonstrate that genomic data provides a more sensitive measure of relatedness. Our analysis reveals a decrease in genetic diversity due to closed breeding, emphasizing the potential for genetic intervention to mitigate negative impact on population fitness. We identify contributing factors to the decreasing genetic diversity including breeding within a closed population, unknown kinship among potential mates, and disproportionate genetic contributions from individual founders. Additionally, we highlight idiopathic myocardial fibrosis (IMF), a common cardiovascular pathology observed in zoologically housed gorillas, and identify a novel genetic variant in the TNNI3K gene that appears to be associated with this condition. These findings underscore the importance of incorporating molecular data into ex-situ population management strategies, and advocate for the adoption of advanced genomic techniques to optimize the genetic health and diversity of zoologically housed western lowland gorillas.

## Introduction

Genetic tools have been used since the origins of zoo breeding programs to maintain genetically diverse *ex-situ* populations. To guide breeding and promote the health of zoological populations, kinship and trait heritability have been determined using pedigree mapping. However, in the application of this method to non-human animal populations, multiple challenges exist including the absence of family structure information for the founding population (incomplete pedigrees), decreased confidence in the paternal contribution (except in controlled breeding conditions), cryptic relatedness, and the inability to differentiate environmental versus genetic contribution to observable traits. The emergence of molecular-based genetic tools including microsatellite mapping, genotyping microarrays, and whole genome sequencing have proven powerful in both scientific and medical applications and have the potential to usefully enhance pedigree-based mapping. Reports incorporating these technologies to manage zoological populations are limited, suggesting an underutilization of these approaches.^1–3^ Advances in genome sequencing have resulted in a significant cost reduction, creating an opportunity to integrate molecular data into zoological population management.

Since the first western lowland gorilla was taken from Africa in 1896, a total of 761 wild-born gorillas have been transferred to zoos or other facilities throughout the world. The capture and transport of wild-living western lowland gorillas (*Gorilla gorilla gorilla*) was prohibited in 1975 under the Convention on International Trade in Endangered Species of Wild Fauna and Flora (CITES). Additionally, the International Union for Conservation of Nature (IUCN) placed the gorilla on the list of critically endangered species and estimates that the wild western lowland gorilla population will reduce by over 80% within the next three generation times (generation time = 22 years).^4^ As a consequence, the maintenance of zoologically housed western lowland gorilla populations relies singularly on captive breeding.

Maintaining genetic diversity within a closed breeding population optimizes adaptive potential and can reduce the prevalence of heritable disease traits. Overrepresentation of disease phenotypes has been reported in zoologically housed western lowland gorillas.^5–9^ The most commonly reported cardiovascular pathology is idiopathic myocardial fibrosis (IMF) also referred to as interstitial myocardial fibrosis and fibrosing cardiomyopathy. Left ventricular hypertrophy and arteriosclerosis of intramural coronary arteries are common accompanying findings with IMF. The pathological presentation has been observed in other great ape species including chimpanzees and bonobos, and shares similarities with the hearts of chronically hypertensive humans, making the investigation of blood pressure one of the priorities in zoologically housed gorillas. Male sex and increased age have been identified as risk factors, and it a complex, mixed etiology has been hypothesized, including dietary factors, obesity, infectious disease, stress, physical inactivity, or genetics.^7,8,10,11^ However, there are no published studies that investigate the familial basis or genetic etiology of IMF.

The zoological western lowland gorilla population was populated exclusively by wild-born individuals until 1956 when the first viable zoologically bred gorilla was born. Zoo breeding was not overwhelmingly successful until the late 1970s when the understanding of life cycle requirements in addition to the manipulation of diet, environmental factors, and medical treatment allowed for greater success in live births.^12^ Since 1956 more than 1,500 western lowland gorillas have been born in captivity. In the 1980’s, as zoological breeding strategies for gorillas became increasingly more successful, both North American and European zoological communities established the respective Gorilla Species Survival Plan® (SSP) and Endangered Species Program (EEP) aimed at ensuring genetic diversity and maintaining demographic health of zoo housed gorilla populations.^13^

Here, we explore the benefits of integrating genomic information for population management using the international zoologic population of western lowland gorillas, focusing on genetic diversity and identifiable heritable diseases. First, we compare kinship values, a measure of relatedness and a proxy for genetic diversity, using classical pedigree mapping and whole genome sequencing-based methods. We then investigate how the origins of this population and other factors have the potential to affect genetic diversity. Lastly, using whole genome sequencing we identify possible genetic determinants of IMF.

## Materials and Methods

### Data

*Studbook*: Information from all zoologically housed gorillas, including studbook number, sex, date of birth, origin (wild born, captive bred), date of death, place of death, cause of death, collection, and relatives, was provided by openly available information from the Dewar Wildlife Gorilla Studbook and the international studbook for the western lowland gorilla.^14,15^ These resources provided information on 2305 western lowland gorillas, including 234 individuals who were stillborn. For the remainder of this paper, we will exclude stillborn individuals from the analysis unless otherwise stated. All data made available from these studbooks were compiled into a comma-separated file (.csv) and used for analysis described throughout the text.

*Whole genome sequencing*: Short read whole genome sequencing data generated by sequencing by synthesis (Illumina inc, San Diego, CA) publicly available on NCBI (accession number: SRP018689) for the western lowland gorillas, Dolly, Abe, Tzambo, and Porta were used for variant discovery.^16^ Variant discovery is described later in the material and methods section.

*Variant call files (VCF):* Files for 26 western lowland gorillas, 3 eastern lowland gorillas (*Gorilla berengei graueri*), and 1 cross river gorilla (*Gorilla gorilla diehli*), were used for kinship analysis and pedigree mapping.^17,18^ Files were generated using high coverage (20-30X) whole genome short read (150bp) sequencing data generated using the sequence by synthesis approach (Illumina inc., San Diego, CA) and aligned to the NCBI35 (hg17) reference genome. This VCF file was used for kinship analysis as described later in the material and methods section.

*Post-mortem reports*: Post-mortem reports were obtained either from the North American SSP for western gorillas, the Great Ape Heart Project or EEP. Non-English written reports were translated, in some instances using Google Translate tools. Idiopathic myocardial fibrosis inclusion criteria required histopathology evidence of fibrotic infiltrates of the myocardium of which no other cause was identified. Interpretation of the postmortem reports was accomplished by the study authors who include licensed veterinarians (JG, HM, VS), a board-certified veterinary pathologist (LL), and board-certified human cardiologists (IK, EA, JP)

### Kinship Analysis and Pedigree Mapping

*Studbook data*: For all 2305 individuals in the western lowland gorilla international studbook, kinships estimates were made for each possible pairing of gorillas using Kinship2 R package.^19^ Using this package, pedigrees were assembled for each family unit, where the founder(s) is/are defined as the first wild born genetic contributor(s).

*Whole genome sequencing data*: From the joint variant call files, where short read (Illumina inc, San Diego) whole genome sequencing data from 30 gorillas was aligned to the NCBI35 (hg17) human reference genome, a bed file was produced using PLINK.^18,20,21^ Using the bed file, kinships were calculated for all possible pairings of these gorillas using KING.^22^

### Network Analysis

A Fruchterman Reingold network analysis was completed with default settings using the Cytoscape tool.^23,24^ Comma-separated files (.csv) were generated that included the gorilla identification number, parentage, country, and zoo of housing.

### Variant Discovery

We aimed to identify individuals with available whole genome sequencing data whose post-mortem reports include detailed histopathology of the myocardium. Individuals whose histopathology report described myocardial fibrosis as moderate or severe, and for which no cause of the fibrosis was otherwise identified, were classified as ‘affected’ and those whose histopathology report showed no evidence of myocardial fibrosis were classified as ‘healthy’. Three (Abe, Porta, and Tzambo) met the affected IMF inclusion criteria and one (Dolly) had no evidence of IMF on histopathology (detailed information for each gorilla is within the supplement). Publicly available fastq files from Illumina short read sequencing were downloaded from NCBI and aligned to both the human (GRCh.38, RefSeq GCF_000001405.39) and Gorilla (Kamillah, RefSeq GCF_008122165.1) reference genomes using the Burrows-Wheeler Aligner (BWA), and variants were called using Genome Analysis Toolkit (GATK) according to GATK best practices.^25,26^ Variant call files were then annotated using Annovar.^27^ Because the genomes were aligned to both the human and gorilla reference genome and the phenotypic status of the gorilla reference is unknown, variants were filtered against the unaffected control gorilla using a custom script written in Python3 and is available on GitHub (https://github.com/jgorz/Great_Ape_Filter.git). Once the VCF was filtered against the unaffected control gorilla, further filtering including identification of variants in genes known to cause cardiomyopathy/cardiovascular (Supplementary Table S2) disease in humans was accomplished using a custom-written bash script available on GitHub (https://github.com/jgorz/Cardiomyopathy_Filter.git).

## Results

### Molecular genomic data is more sensitive at identifying relatedness than historical pedigree data

To investigate the reliability of the studbook in predicting relatedness among paired individuals, we compared pedigree-derived kinship values (**Supplementary Table 1a**) to whole genome sequence-derived kinship values (**Supplementary Table 1b**) from 435 unique pairings of 3 captive eastern lowland gorillas, 1 captive cross river gorilla, 9 zoo born western lowland gorillas and 17 wild captured western lowland gorillas (**Figure 1**). Pedigrees derived from the international studbook suggested 9 pairings to share relatedness (range 2-5th degree relatives). WGS-derived kinship values confirmed the 9 pairs to be related and identified an additional 5 related pairs (**Table 1**) revealing the sensitivity of the studbook to identify related pairs to be 64.28%.

**Figure 1.**
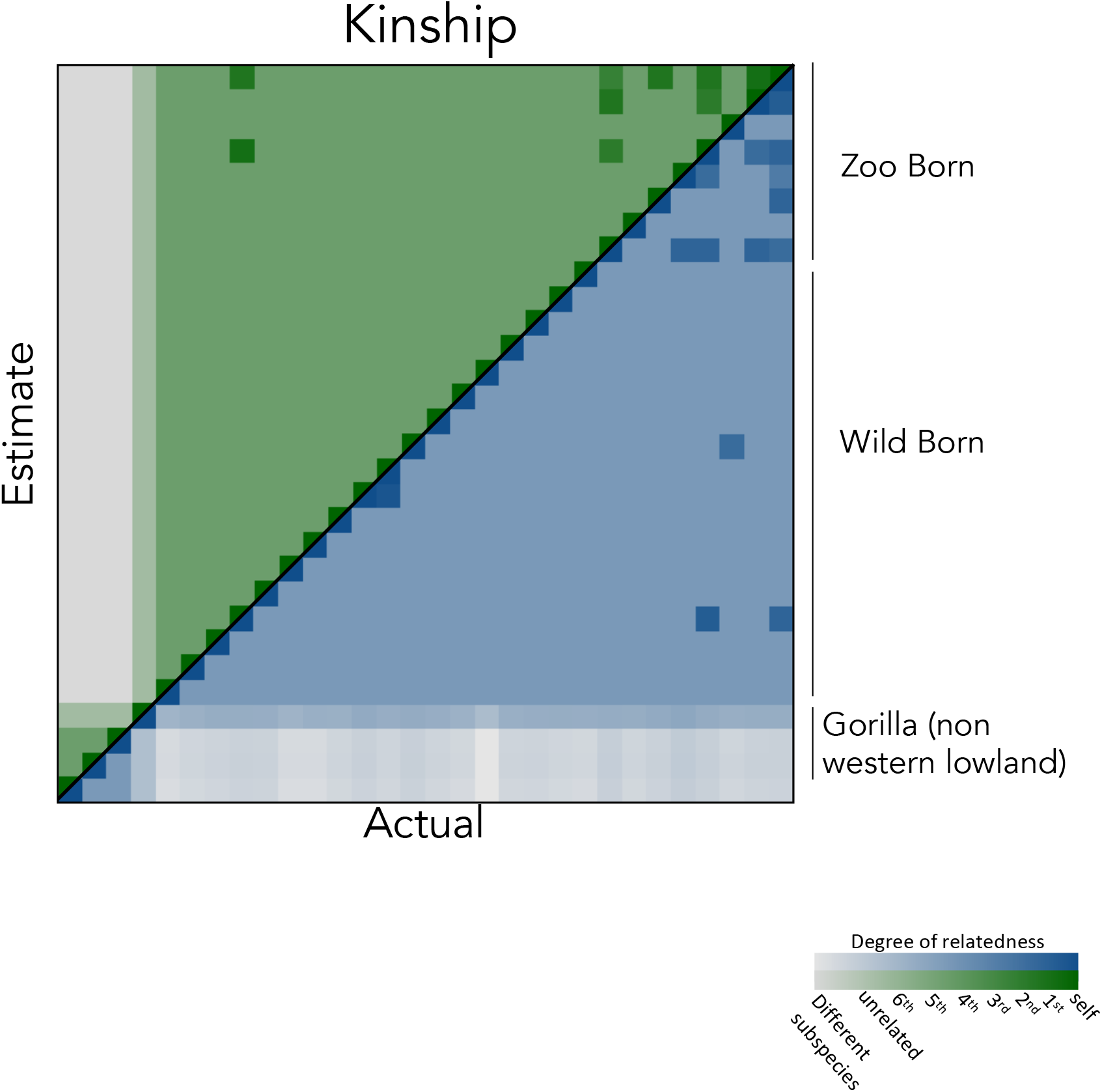
Pairwise comparison of estimated (pedigree derived) kinship to actual (whole genome sequence derived) kinship reveals discordance among 5 western lowland gorillas. (Green) estimate kinship from pedigree obtained from the international studbook. (Blue) Actual kinship calculated from whole genome sequencing data using the PLINK tool KING. Each individual box represents the kinship of two individuals. Darker colors indicate higher kinships.

**Table 1:** Comparison of kinship and degree of relatedness values calculated from pedigree data or whole genome sequencing (WGS) data reveal discordance. Of the 14 pairings that revealed relatedness using whole genome sequencing data, 9 were predicted using pedigree data and 14 were observed using WGS data. WB=wild born, ZB=zoo born. Relatedness=Degree of relation.

| ID 1 | ID 2 | Estimate (pedigree) |  | Actual (WGS) |  |
| --- | --- | --- | --- | --- | --- |
|  |  | Kinship | Relatedness | Kinship | Relatedness |
| WB-001 | WB-002 | 0 | 0 | 0.2268 | 1 |
| ZB-001 | ZB-002 | 0 | 0 | 0.0504 | 3 |
| ZB-003 | ZB-002 | 0 | 0 | 0.024 | 4 |
| WB-003 | ZB-004 | 0 | 0 | 0.0214 | 4 |
| ZB-005 | ZB-002 | 0 | 0 | 0.0032 | 6 |
| ZB-006 | ZB-005 | 0.125 | 2 | 0.141 | 2 |
| ZB-007 | ZB-003 | 0.125 | 2 | 0.1124 | 2 |
| ZB-001 | ZB-006 | 0.078 | 3 | 0.0636 | 3 |
| ZB-003 | ZB-005 | 0.0625 | 3 | 0.0482 | 3 |
| ZB-007 | ZB-005 | 0.0625 | 3 | 0.0465 | 3 |
| ZB-008 | ZB-005 | 0.0625 | 3 | 0.0398 | 3 |
| ZB-001 | ZB-003 | 0.0313 | 4 | 0.0573 | 3 |
| ZB-006 | ZB-003 | 0.0313 | 4 | 0.0205 | 4 |
| ZB-001 | ZB-005 | 0.0156 | 5 | 0.0204 | 5 |

The unexpected kinship reveals inaccuracies within the pedigrees derived from the studbook. The observed first-degree relatedness between WC-001 and WC-002 revealed by genomic data directly counters the assumption that wild-captured gorillas share no relatedness. Additionally, the unexpected relatedness shared between the other pairings suggests that these individuals are descendants of a common wild ancestor, further illustrating the weakness in this assumption. According to this data, kinship calculated from whole genome sequence data is a better predictor than pedigree data.

To investigate the reliability of the studbook as a predictor of genetic diversity, we compared the kinship of 325 unique pairings from 26 western lowland gorillas using either *non-genetic* pedigree data from the studbook or *genetic* whole genome sequencing data. Kinship values from non-genetic and genetic data revealed a mean of 0.0018 (average of > 7th-degree relatives) and 0.0027 (average of > 8th-degree relatives) (p = 0.23) (**Table 2a**). Although the means are not significantly different, the use of genetic data uncovered a higher mean kinship, suggesting that non-genetic data may not be a reliable predictor of genetic diversity within the zoological western lowland gorilla population.

**Table 2.**
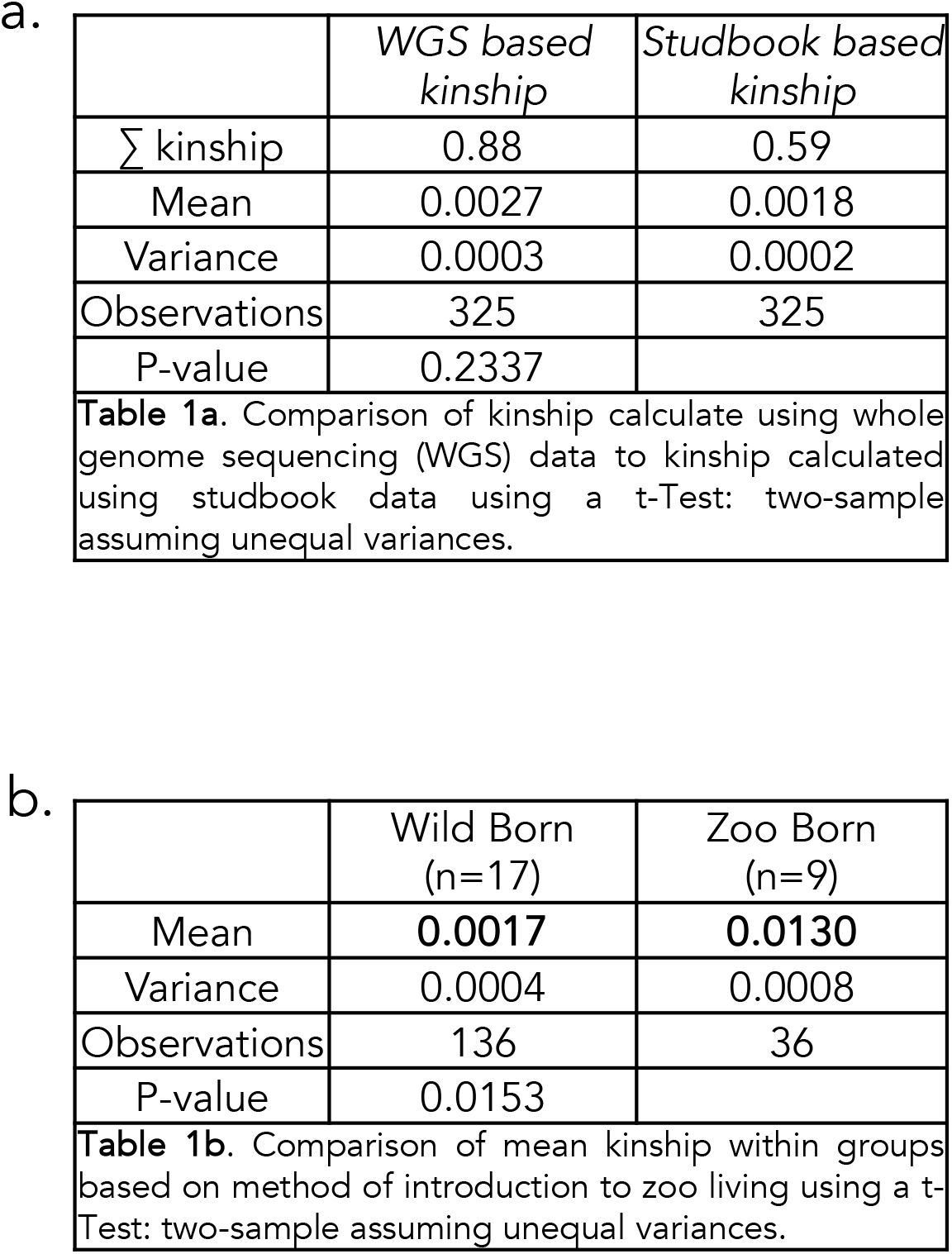

### Whole genome sequencing data identifies decreasing genetic diversity as a result of a closed breeding population and other factors

#### Origin of the Breeding Population

The current population of zoologically housed western lowland gorillas is comprised of individuals who were either taken from their native habitats in central Africa (wild-born) or were bred and born in zoos (zoo-born). When we investigate the chronology of the two introduction methods among 760 wild-born and 1310 zoo-born gorillas, we identify a bimodal distribution (**Figure 2a**) with a peak of wild-born introduction in 1970. Of the 760 wild-born gorillas 100 males and 144 females mated successfully in zoos, giving rise to the first generation of zoo-born gorillas (**Figure 2b**) designating these 244 wild-born gorillas as the founding population.

**Figure 2.**
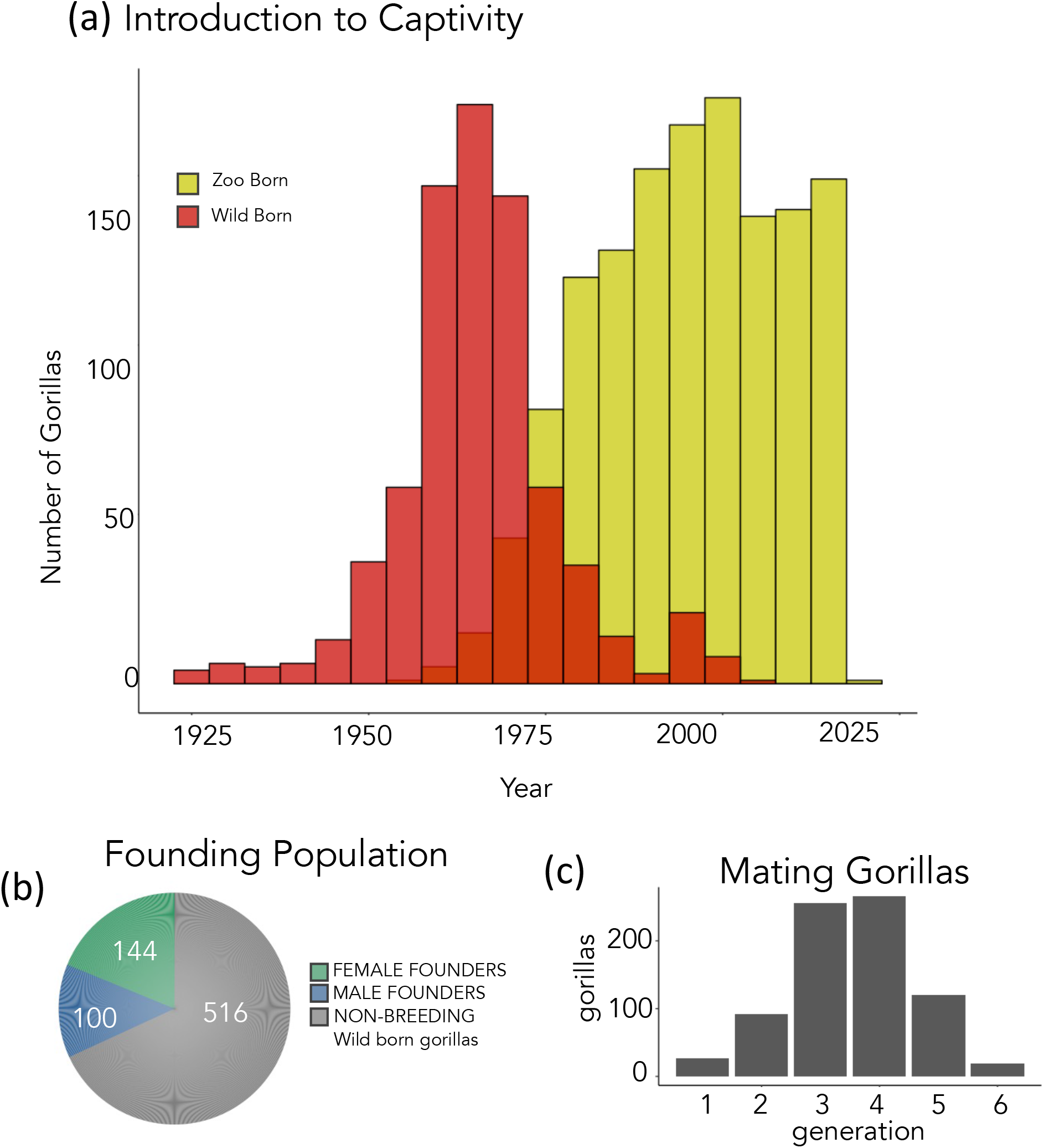
The current breeding population is composed of zoo born individuals originating from 100 male and 144 female founders. (a)Timeline of gorilla introduction to captivity: number of individuals introduced to captivity grouped by method of introduction (Red-wild born gorillas, Yellow-zoo born gorillas) in five year intervals. (b) Founding Population: Wild born gorillas who have genetic contributions to the zoo born population (founders) based on on sex (green-Female founders, blue-male founders). (c) Distribution of generation time of gorillas with mating potential (less than 35 years).

Genetic contribution from individuals outside of zoo populations has been restricted, and as a result, maintenance of the zoo-housed western lowland gorilla populations is reliant on breeding individuals from within this population. We expect to observe a loss of genetic diversity as the generation time of breeding individuals increases within a closed population. We therefore investigate the origin and generation times of western lowland gorillas with the potential to breed, defined as any individual under the age of 35 years. 739 gorillas are suspected to be fertile, and 719 of these are descendants from the zoological founding population, with a generation time ranging from 2-6 (**Figure 2c**). The remaining 20 gorillas are wild born (1st generation). Of these, 15 have not successfully mated and 13 are geographically isolated, reducing the chances of them contributing to the global zoological western lowland gorilla gene pool. This data shows that the gene pool is primarily restricted to the 719 individuals with close relation to 244 founders. As breeding continues within the closed population we expect to see a decrease in genetic diversity.

#### WGS: Closed breeding decreased genetic diversity

Using whole genome sequencing data, we investigated how closed breeding may affect the genetic diversity of the zoo-housed western lowland gorilla population. To identify how this affects genetic diversity, we grouped individuals based on their origin (wild born-wild born (WB-WB), zoo born-zoo born (ZB-ZB)) and calculated the individual mean kinship (average kinship shared between an individual and all others within the group) values from 26 zoo housed individuals and compared the group mean kinship from WB-WB and ZB-ZB groups. Of the WB-WB group we found the population mean kinship to be 0.0016 suggesting that the wild-born population shares minimal relatedness. Among the zoo-born population, we found the mean kinship to be 0.0114 suggesting that on average zoo-born individuals are separated by five meiotic events. A statistically significant difference between the two groups was observed (p-value=0.013) (**Table 2b**), suggesting that as breeding in the closed population continues and the generation time of breeding individuals increases, genetic diversity will subsequently decrease.

### Founder contributions to the current breeding population

Sexual selection associated with hierarchical social structuring within wild gorilla troops in native habitats likely lead to an abnormal distribution of founder contributions to the gene pool. However, in zoos, mate selection is primarily a result of human management, through breeding recommendations, management of troop structures, and use of contraceptives. Managed carefully, this could result in an evenly distributed founder contribution to the current gene pool and ease the management of the zoological breeding population.

To investigate this, we calculated the genetic contribution of each founder to the current breeding population gene pool. (Summation of the kinship coefficients calculated between a single founder and all breeding individuals divided by the summation of kinship coefficients calculated between all founders and breeding individuals, **Figure 3a**). Of the 100 male and 144 female founders, 86 and 114 have direct descendants in the current breeding population, respectively. 3.05, 2.97, and 2.86% of the entire breeding gene pool were contributed by three male founders, and 2.65, 2.15, and 2.10% were contributed by three females, totaling 15.78%. When we investigate the generation number of the direct descendants of these ‘major contributors,’ we observe that the majority are 3rd and 4th generation, meaning 2 to 3 meiotic events separate the majority of the direct descendants from their founding ancestor (**Figure 3b**). We further calculated the number of individuals within the current breeding population with genetic contributions from each founder. Of the 780 individuals, 13.8, 20.6 and 10.6% have genetic contributions from the top three male founders, respectively, and 11.4, 17.7 and 10.3% have genetic contributions from the top three female founders, respectively (**Figure 4a**). In total, 51.3% of the breeding population have genetic contributions from at least one of the top three male or female genetic contributors (**Figure 4b**). (Pedigrees have been mapped for each of the top genetic contributors (**Supplementary** Figures 1e-j))

**Figure 3.**
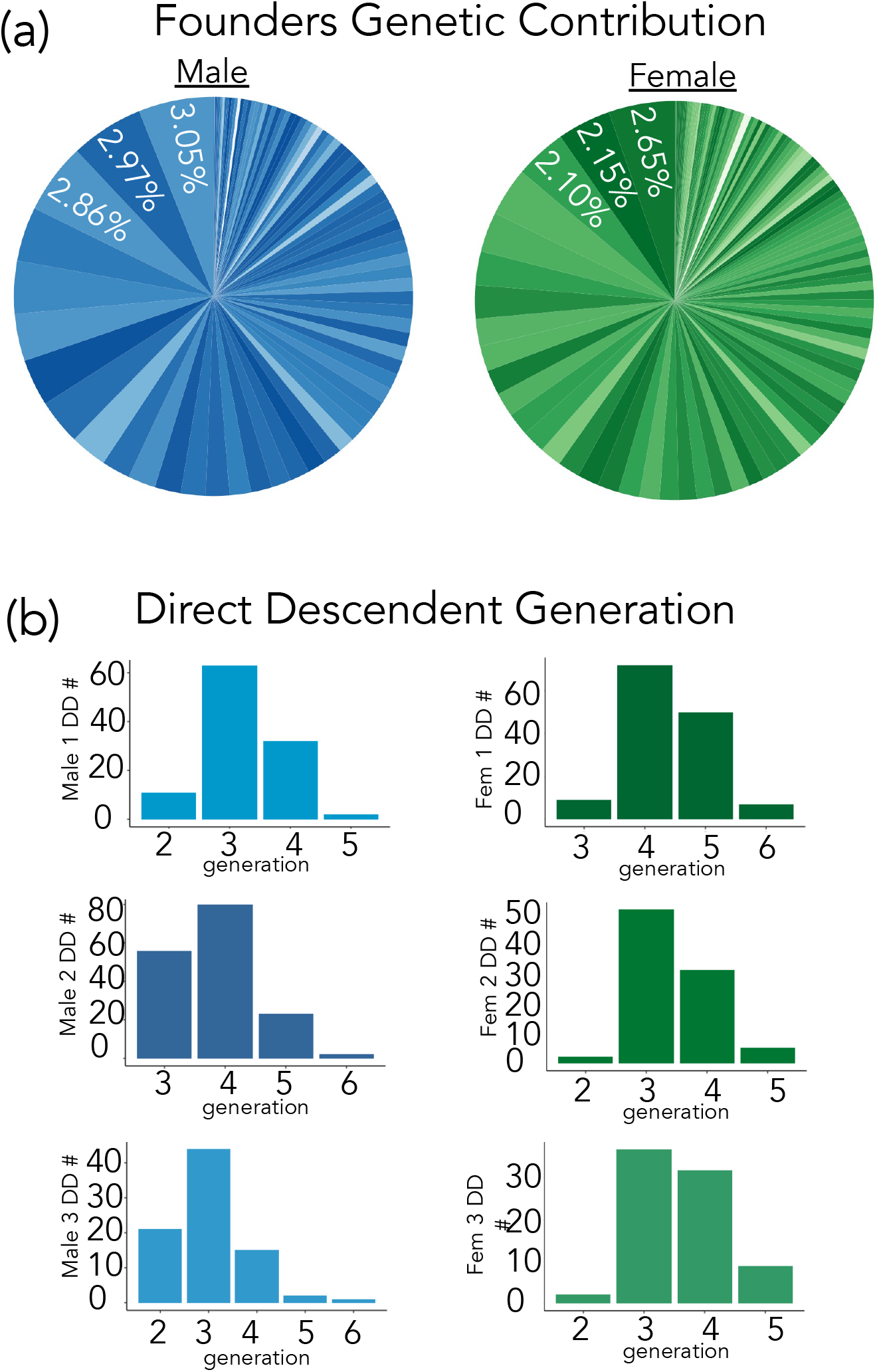
A large proportion of the current breeding gene pool was contributed by a small group of founders. **(a)** Percentage of all individual founders genetic contribution (blue pie chart=male founders, green pie chart=female founders. **(b)** Distribution of the generation number of each of the direct descendants from the founders with the top genetics contributions. Generation 1 is designated as the wild born founders. Generation >1 are zoo born individuals.

**Figure 4.**
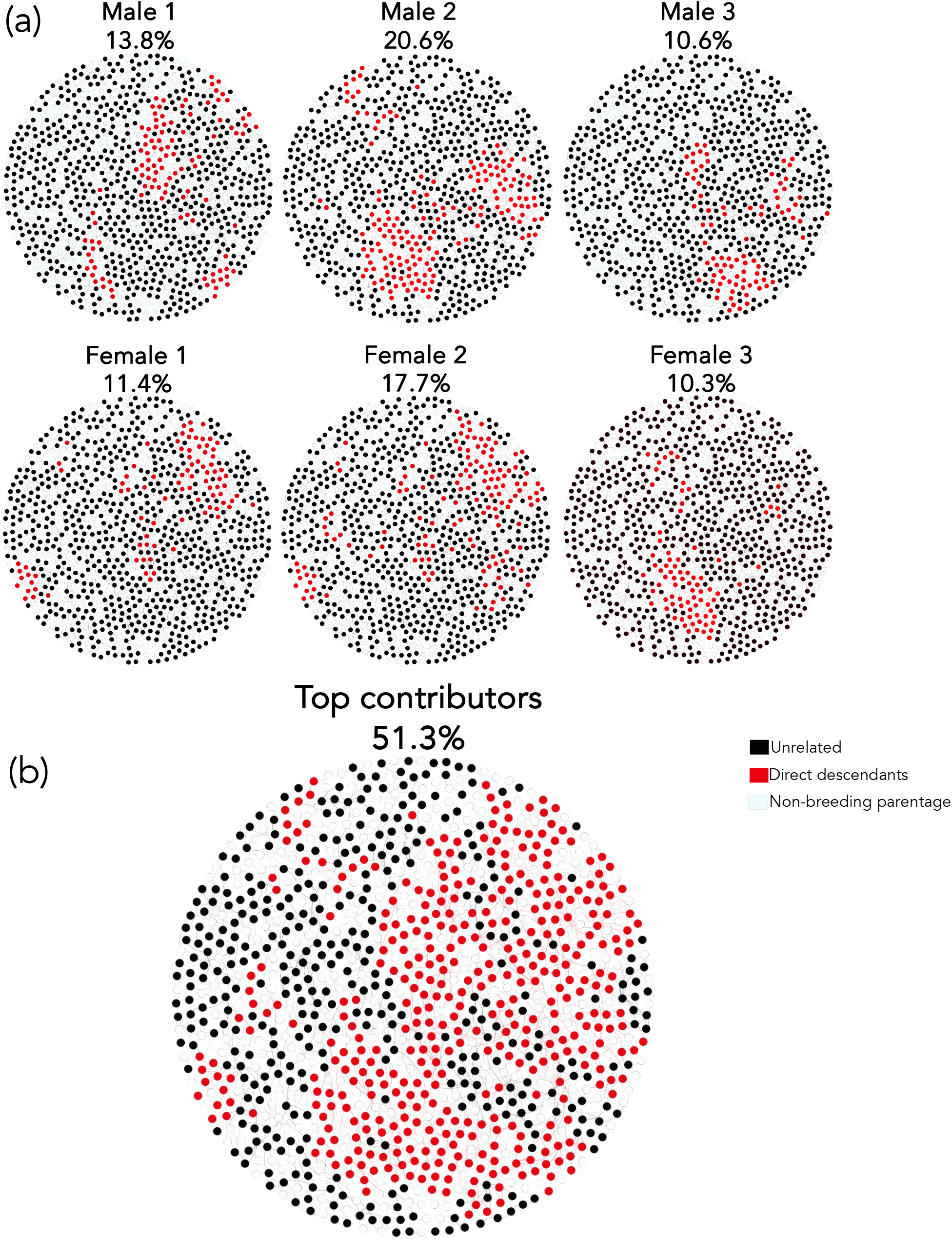
Large proportion of the current breeding population are direct descendants of the top genetic contributors. Using a Fruchterman Reingold network analysis of the breeding population where nodes are individual gorillas and edges as parentage. **(a)** Individuals with lineage to the top three male and female genetic contributors to the gene pool. **(b)** Individuals related to at least one of the top male or female contributors. (white nodes indicate parents who no longer have the ability to mate due to death or advanced age)

### Geographical Bottlenecks

Optimizing mate selection is the first-line strategy for maintaining a genetically diverse population. However, geographical isolation is a limiting factor for mate selection and may create bottlenecks within zoologically-housed gorilla populations. To investigate the role of geography in the mating patterns of gorillas under human care, we performed a Fruchterman Reingold network analysis of the entire zoo gorilla population designating nodes as individual gorillas (**Figure 5**).^24^ Although we observe the majority of breeding pairs are from within the same zoo, we find that transport of individuals for breeding purposes within North America or Europe is prevalent. However, the transport of individuals for breeding purposes between North America and Europe is limited, resulting in two largely isolated breeding populations. We identified one individual who contributed to the gene pool in both North American and European zoo populations. This individual was captured from Africa in 1961 and was transported to North America where he produced two offspring between 1970-1971. He (184) then was transported to Europe in 1973 where he produced 12 offspring between 1975 and 1982. He returned to North America where he lived the rest of his life with no further reproduction. Of the 12 offspring born in Europe, one was transported to North America where he successfully bred multiple times. We further identified 7 (573, 848, 749, 682, 885, 752, 687) individuals who were born in Europe to European-housed parents and were then transported to the US where they bred successfully. These were the only events where successful breeding was accomplished after individuals were shared between European and North American countries. Continued geographical isolation will increase the loss of genetic diversity.

**Figure 5.**
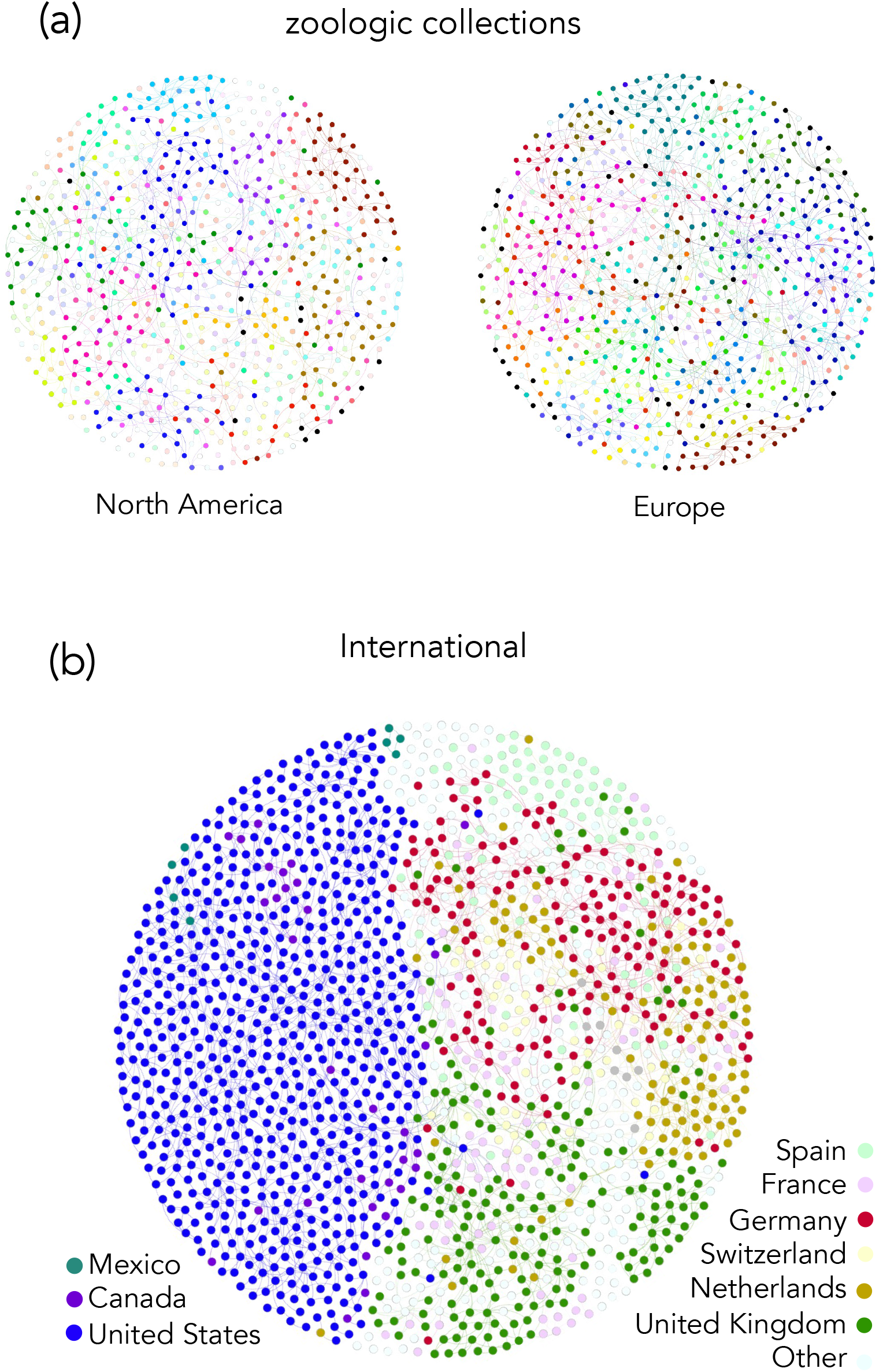
Fruchterman Reingold force directed graph demonstrates a bottleneck across the atlantic ocean: Nodes: individual gorillas colored by location of housing, Edges: parentage. **(a)** Shows individuals housed in north American or European zoos and are colored based on the zoologic population they are from. **(b)** Shows the international gorilla population and are colored based on country.

### Risk of undesirable heritable traits

We have investigated four factors that have the potential to reduce the genetic diversity within the zoologically housed western lowland gorilla population. Lack of genetic diversity will reduce the adaptive potential of a population and will increase the prevalence of traits inherited from the founder population. Cardiovascular disease has been widely reported to have a high mortality rate in zoo-housed great apes, with the most commonly reported phenotype described as idiopathic myocardial fibrosis.^5–8^ Where available, using health records and post-mortem reports in combination with the studbook we look for aggregation of disease within founder pedigrees. Here we present two pedigrees from male founders who were housed in North American Zoos (**Figure 6**). The founding individual from pedigree 1 was wild born and introduced to captivity in 1969 and died in 2015 at the age of 46 years. He was observed to have myocardial fibrosis at post-mortem. Among his ten offspring (8 Males, and 2 Females, born between 1986-2007), 1 male died with a similar disease phenotype at the age of 18 years (1993-2011), and three additional offspring have had myocardial structural and functional abnormalities observed during routine health screening and echocardiograms. The founding individual from pedigree 2 was a wild born gorilla introduced to captivity in 1962 who died in 2005 at the age of 43 years due to complications of sepsis. For this individual no cardiac abnormalities were noted during the necropsy. This founder had 7 offspring (2 Males, and 5 females), and none has shown signs of cardiac abnormalities. Phenotypic aggregation within familial populations exemplified by these two pedigrees suggests that idiopathic myocardial fibrosis may be a heritable trait but data is limited to allow definitive conclusions.

**Figure 6:**
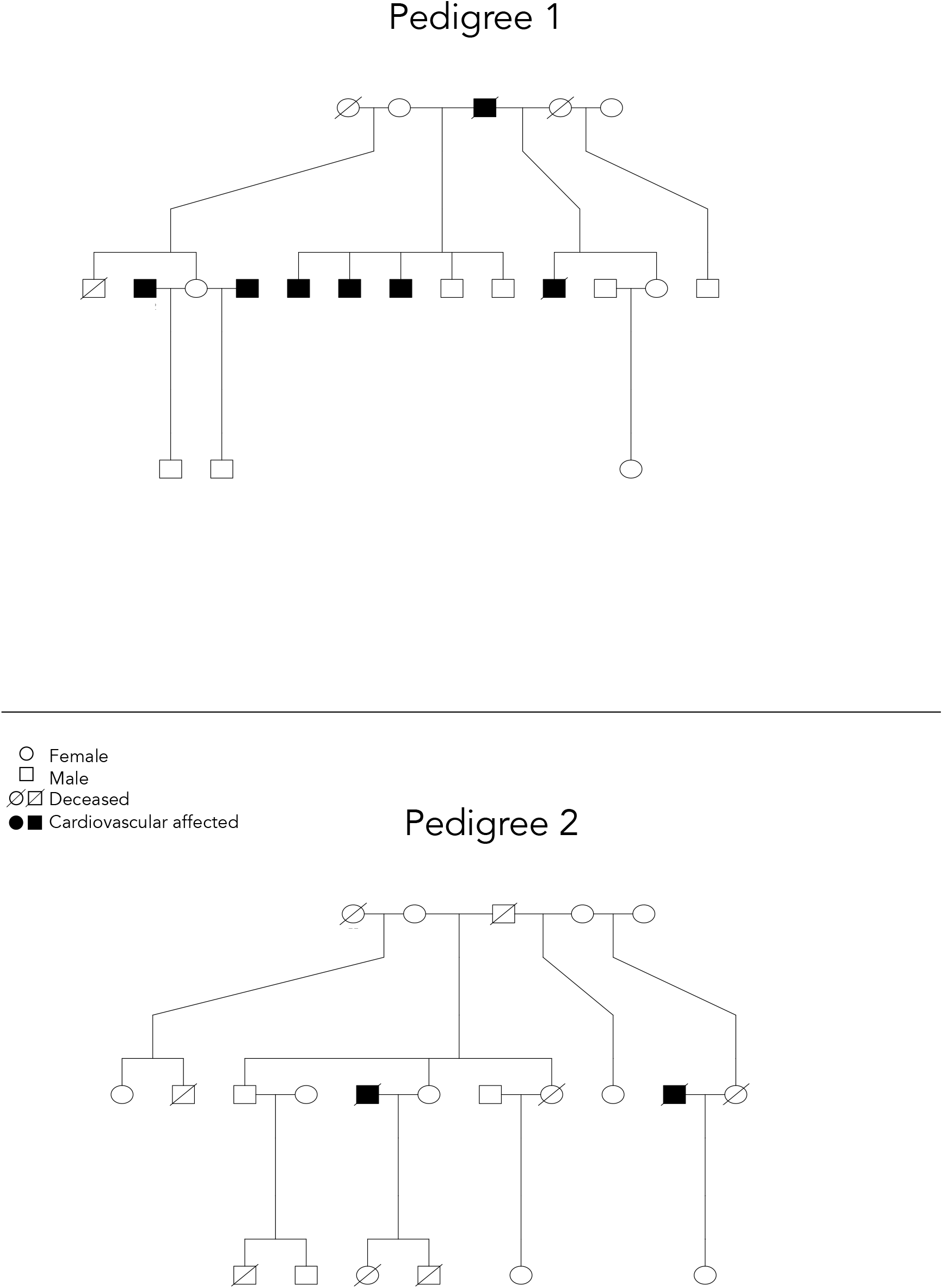
Pedigree mapping provides evidence of familial aggregation. Four direct descendants of the affected male founder from Pedigree 1 have evidence of cardiovascular disease. Zero direct descendants of the unaffected male found from Pedigree 2 have evidence of cardiovascular disease.

Of the 200 founders contributing to the current breeding population, 150 are deceased. Of the deceased individuals, post-mortem reports are available for 65 individuals. Of these, 35 are reported to have notable cardiac pathology, and 32 (49%) of these individuals have evidence of myocardial fibrosis. Among the three male and three female top genetic contributors (Figures 2+3), one male is living with no medical records available, two males are deceased with postmortem reports available, and no reports are available for the three females. Of the two males with an available postmortem report, one died of bacterial meningitis with no reporting of myocardial fibrosis at the age of 32, and the other (male #2) died of an aortic dissection with severe myocardial fibrosis observed on myocardial histopathology at the age of 25. 22.1% of the zoologically housed population with the potential to breed are direct descendants of male #2. If genetic determinants are associated with idiopathic myocardial fibrosis, then the current population and future generations of zoo-born gorillas may experience an increased prevalence of this pathology phenotype making a strong case for molecular investigation of possible causes.

### Identification of genetic determinants of IMF

To investigate a genetic etiology of idiopathic myocardial fibrosis, we identified four unrelated individuals who had both myocardial histopathology performed during necropsy and available whole genome sequencing data. Of these four individuals (2 male (m), 2 female (f)); Dolly’s (f) myocardium was reported to be healthy with no evidence of fibrosis, and the other three individuals (Abe (m), Porta(f), Tzambo(m)) were reported to have myocardial fibrosis. Their genomes were aligned to the human reference HG38 and variants were called using GATK best practices. Variants were then filtered according to the schema (**Figure 6**). In a total 97 exonic variants segregated to affected individuals in genes known to be associated with human cardiovascular disease (**Supplementary Table 2**). These variants were then prioritized based on *in-silico* variant predictors, SIFT, and Polyphen (full list of variants available in the supplement).^28,29^ A missense variant in exon 23 of *TNNI3K* (c.C2182A p.L728M) the gene that codes for cardiac troponin-I interacting kinase, resulting in a lysine to methionine replacement at amino acid position 728 was ranked with highest priority. This novel variant has not been reported in humans or gorillas.^30^ Variants identified in TNNI3K in humans have been associated with supraventricular tachycardia, conduction disease and cardiomyopathy making this a strong candidate for causality. ^31–34^

## Discussion

The integration of genomic data into the management of zoologically housed western lowland gorillas represents a pivotal step in ensuring the genetic diversity and health of this endangered species. While prior studies have acknowledged the efficacy of microsatellite testing for assessing ancestral genetic population structure and relatedness among breeding pairs, and have established best practices for molecular data utilization in population management, its application in zoological populations has been underutilized.^1,3^ Our research not only supports these findings but also emphasizes the need to shift from conventional pedigree-based methods towards more sensitive molecular-based approaches for evaluating relatedness and genetic diversity. Moreover, our utilization of whole genome sequencing has revealed potential heritability of idiopathic myocardial fibrosis, demanding further inquiry into its genetic underpinnings.

Using whole genome sequencing data our study has revealed a decline in genetic diversity among the zoologically housed western lowland gorilla population. We have identified several factors contributing to this decline including breeding within a closed population, unknown kinship among potential mates, and significant genetic contributions from individual founders. The finding that over 50% of the current breeding population are direct descends of just six founders is a striking observation. The overwhelming genetic contribution of these six founders diminishes the pool of suitable mating pairs, underscoring the necessity for accurate determination of kinship between potential mates.

Furthermore, our study has illuminated that relatedness estimates derived from classical pedigree mapping lack the sensitivity of those calculated from whole genome sequencing data. We have identified instances of cryptic relatedness among five pairs of zoologically housed individuals. This finding can be explained by incomplete pedigree mapping resulting from a lack of comprehensive family structure data when wild individuals were introduced into zoos. To sustain genetic diversity and the overall health of zoological populations, cooperative agreements have been established among accredited institutions, adhering to the breeding recommendations set forth by the Species Survival Plan (SSP) for Gorillas. These guidelines advocate for mate selection that takes into account factors such as relatedness, reproductive age, and behavioral and health traits.^35^

While leveraging molecular data for mate selection holds the potential to preserve adequate genetic diversity in zoological populations for generations, achieving successful pairings may be challenging. Geographical constraints and the associated risks of transporting gorillas may prevent the introduction of optimal pairs. Additionally, the autonomous nature of gorillas, as highly complex and social animals, introduces a potential barrier to successful reproduction when individuals are introduced into new habitats and social structures.

Assisted reproductive techniques, encompassing gamete preservation, in vitro fertilization, and artificial insemination, offer potential solutions to these inherent and logistical challenges. While reports of these techniques are limited, success rates remain low.^36–38^ Funding and further research into assisted reproductive techniques may prove instrumental in upholding genetic diversity and the overall health of ex situ western lowland gorilla populations.

While the use of whole genome sequencing data holds promise for enhancing genetic diversity and health in ex situ populations, there is a need for accessible expertise in bioinformatics and genetics. To address this, we’re ready to provide a user-friendly pipeline for analyzing raw whole genome sequencing data in Fastq format. This tool will assist in determining kinship and identifying potential genetic factors influencing observable traits. Additionally, we encourage the use of resources available through the AZA Molecular Data for Population Management Scientific Advisory Group.

**Figure 7:**
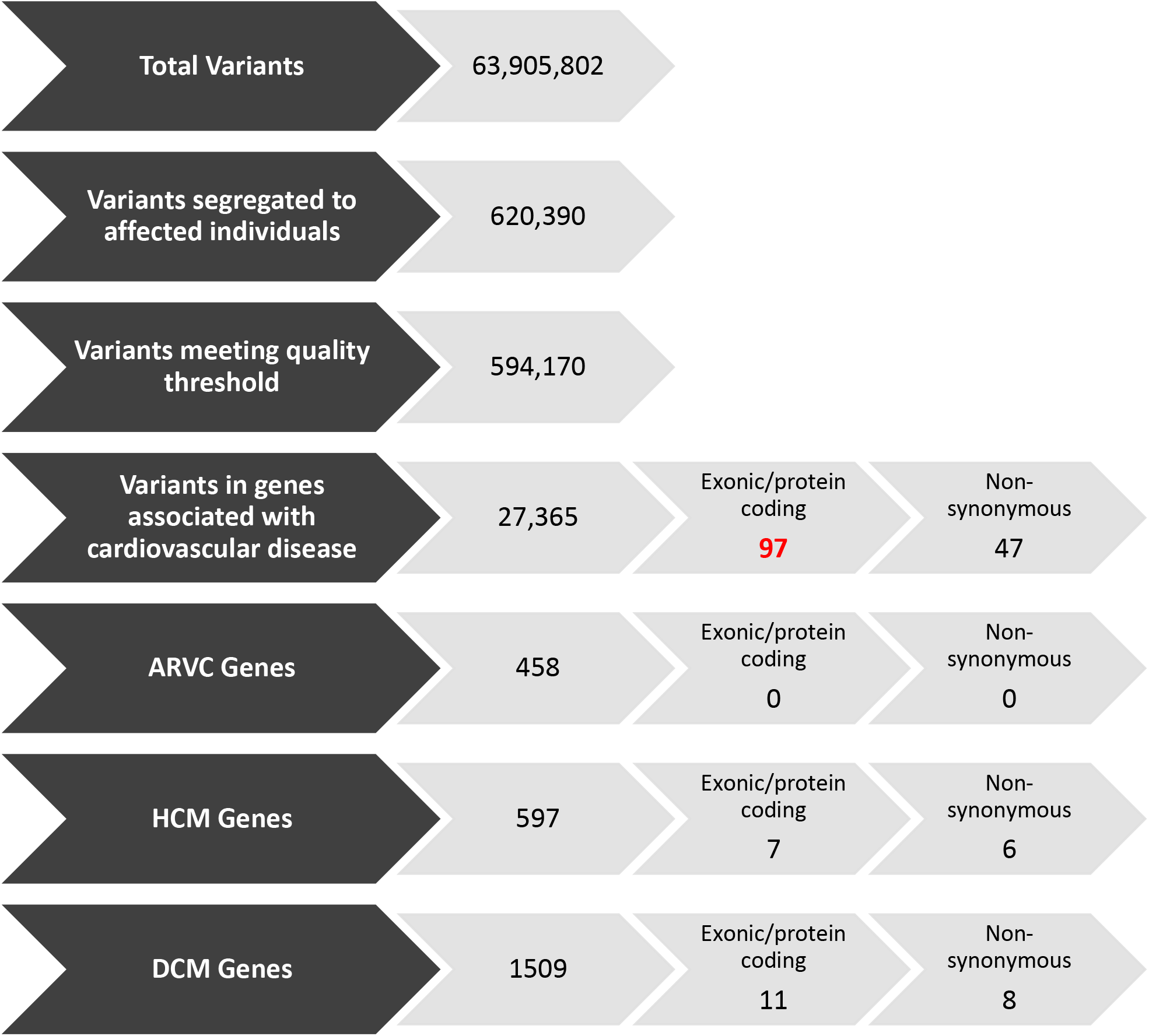
Variant filtering schema. Whole genome sequencing data from three affected and one unaffected individual was aligned to the human reference genome (hg38) and variants were called. The first filtering criteria was the presence of a variant in all three affected individuals (Abe, Porta, and Tzambo) and absent in the unaffected individual (Dolly). Variants were then filtered based on quality score and prioritized if identified in genes known to be associated with cardiovascular disease or cardiomyopathy (ARVC = Arrhythmogenic Right Ventricular Cardiomyopathy, HCM = Hypertrophic Cardiomyopathy, DCM = Dilated Cardiomyopathy). All variants that were then identified in exonic regions were manually curated (red text) with higher priority given to non-synonymous variants.

**Table 3:** Top variant candidates after filtering and prioritization. SIFT value < 0.05 = ‘Deleterious’. Polyphen Value > 0.908 = ‘Probably Damaging’.

| Variant Information |  |  |  | Effect Prediction |  |  | Genotype |  |  |  |
| --- | --- | --- | --- | --- | --- | --- | --- | --- | --- | --- |
| Chr | Gene | Type | Location | SIFT | Polyphen HDIV | Polyphen HVAR | Dolly (-) | Abe (+) | Porta (+) | Tzambo (+) |
| 1 | *TNNI3K* | missense | Exon 23<br>c.C2182A<br>p.L728M | 0.030 | 0.999 | 0.952 | 0/0 | 1/1 | 1/1 | 1/0 |
| 14 | *SYNE2* | missense | Exon 46<br>c.C7259A<br>p.S2420Y | 0.005 | 0.942 | 0.729 | 0/0 | 1/1 | 1/0 | 1/1 |
| 17 | *KCNJ12* | missense | Exon 3<br>c.C1036T<br>p.H346Y | 0.001 | 0.999 | 0.932 | 0/0 | 1/0 | 1/0 | 1/0 |
| 17 | *ABCA10* | missense | Exon 7 c.C528A<br>p.F176L | 0.002 | 1.000 | 0.998 | 0/0 | 1/0 | 1/1 | 1/0 |
| 5 | *VCAN* | missense | Exon 7<br>c.G4122C<br>p.Q1374H | 0.001 | 0.995 | 0.874 | 0/0 | 1/0 | 1/0 | 1/0 |
| 9 | *ANKRD18A* | missense | Exon 10<br>c.A1978G<br>p.K660E | 0.004 | 0.880 | 0.899 | 0/0 | 1/0 | 1/0 | 1/0 |

