## Supplementary Tables 1a, 1b, 2, Figures 1a-j for "Leveraging whole genome sequencing to promote genetic diversity and population health in zoo-housed western lowland gorillas"

### Slide 1
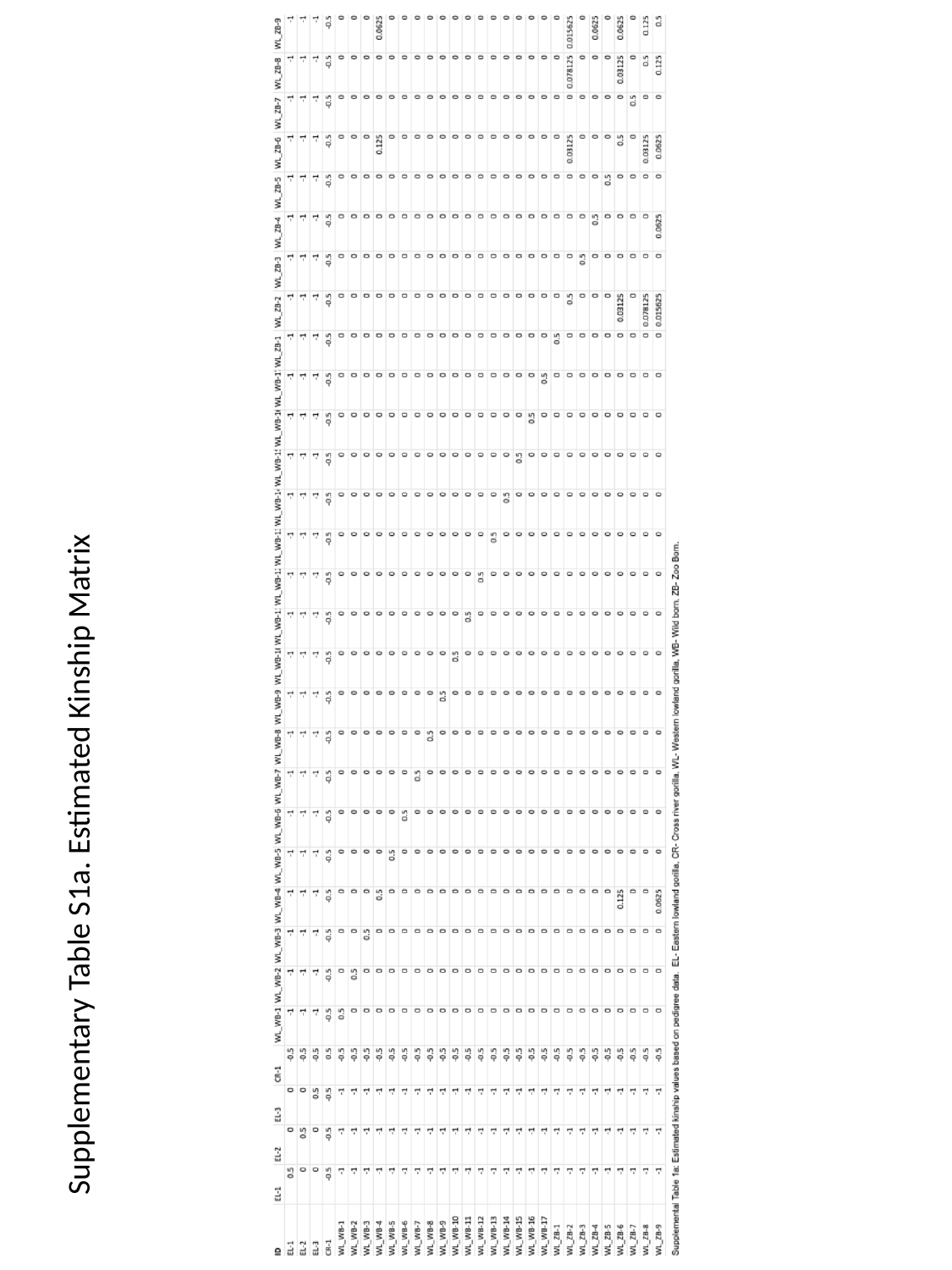

Supplementary Table S1a. Estimated Kinship Matrix

### Slide 2
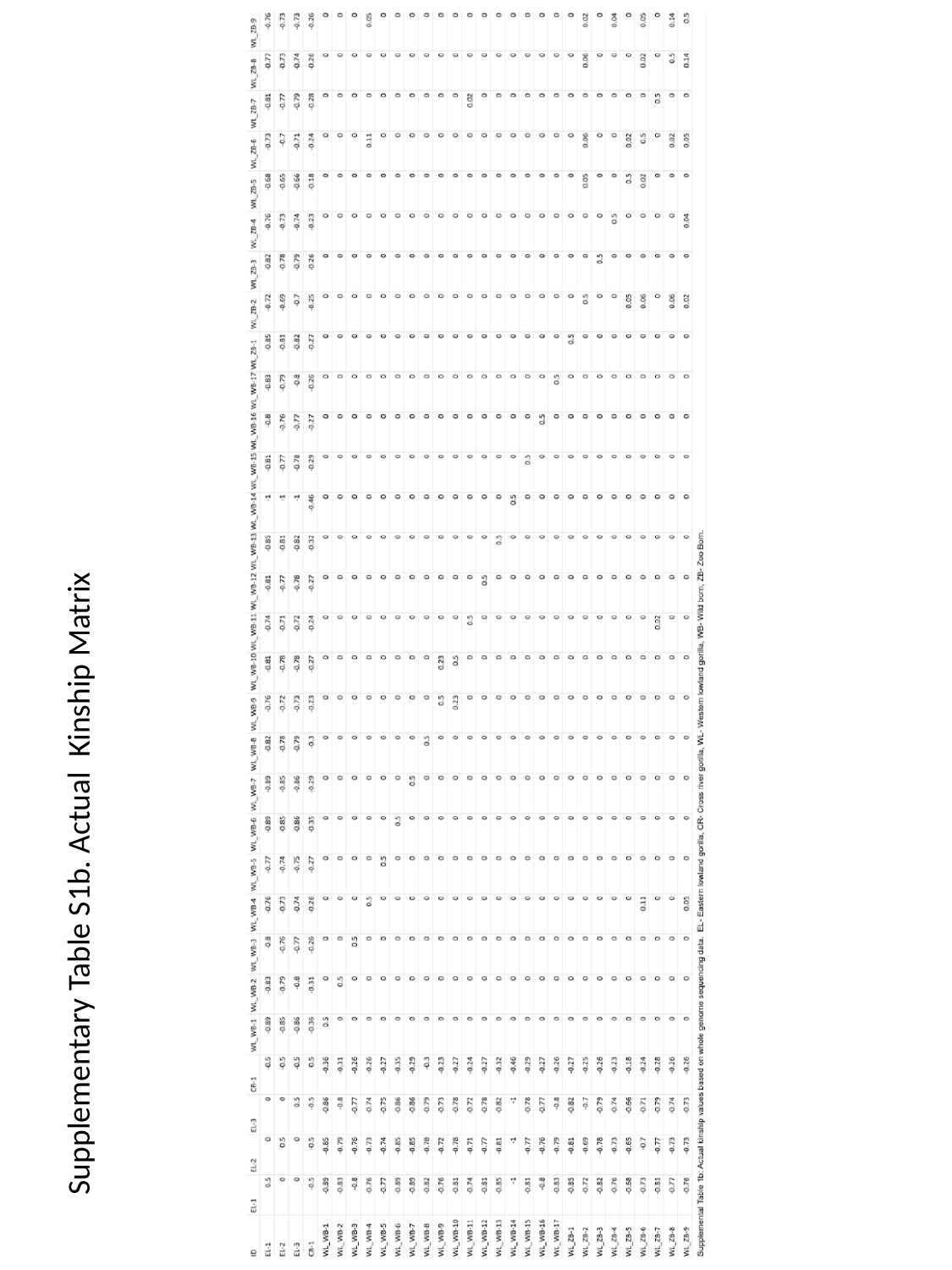

Supplementary Table S1b. Actual Kinship Matrix

### Slide 3
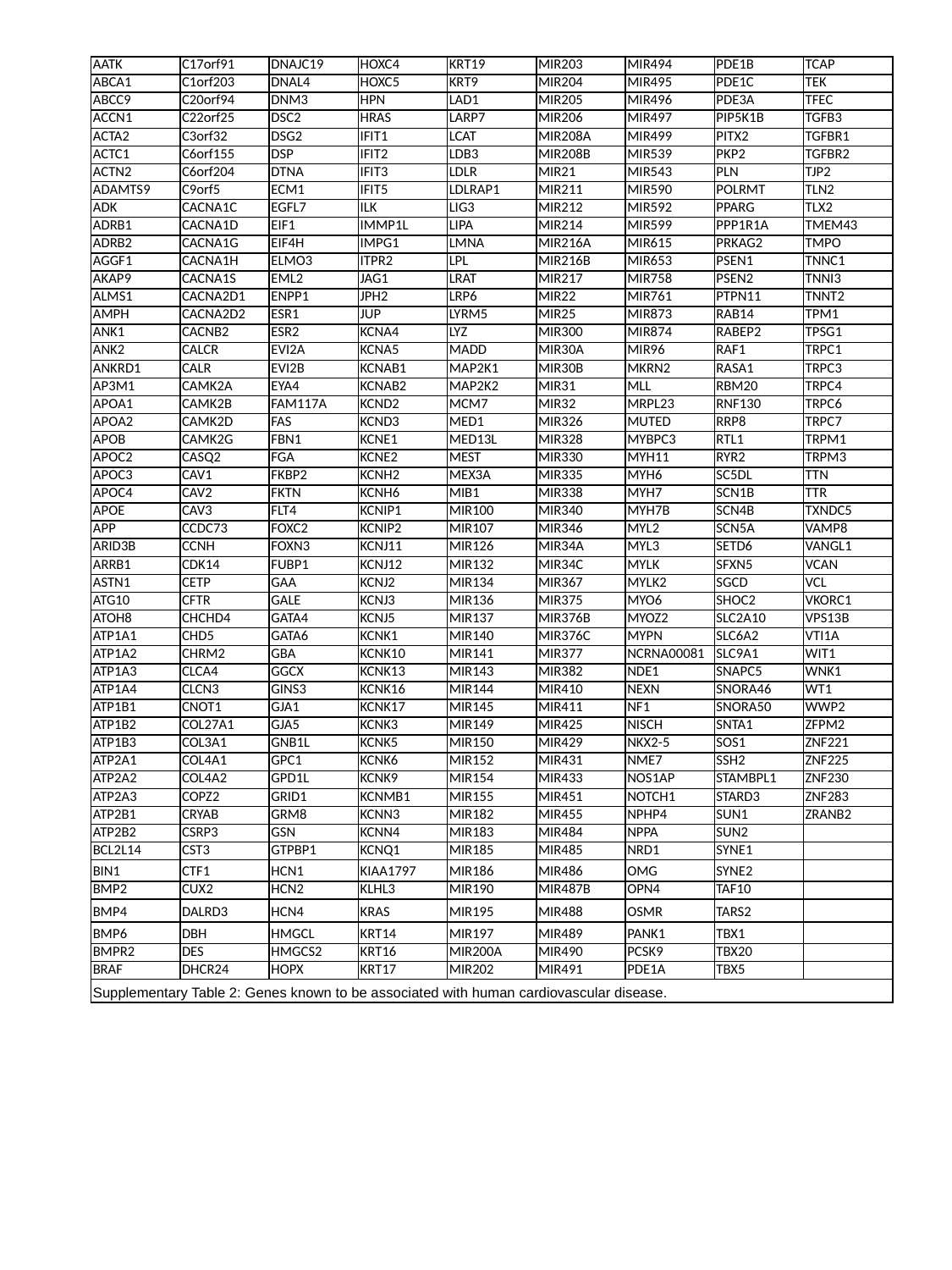

| AATK | C17orf91 | DNAJC19 | HOXC4 | KRT19 | MIR203 | MIR494 | PDE1B | TCAP |
| --- | --- | --- | --- | --- | --- | --- | --- | --- |
| ABCA1 | C1orf203 | DNAL4 | HOXC5 | KRT9 | MIR204 | MIR495 | PDE1C | TEK |
| ABCC9 | C20orf94 | DNM3 | HPN | LAD1 | MIR205 | MIR496 | PDE3A | TFEC |
| ACCN1 | C22orf25 | DSC2 | HRAS | LARP7 | MIR206 | MIR497 | PIP5K1B | TGFB3 |
| ACTA2 | C3orf32 | DSG2 | IFIT1 | LCAT | MIR208A | MIR499 | PITX2 | TGFBR1 |
| ACTC1 | C6orf155 | DSP | IFIT2 | LDB3 | MIR208B | MIR539 | PKP2 | TGFBR2 |
| ACTN2 | C6orf204 | DTNA | IFIT3 | LDLR | MIR21 | MIR543 | PLN | TJP2 |
| ADAMTS9 | C9orf5 | ECM1 | IFIT5 | LDLRAP1 | MIR211 | MIR590 | POLRMT | TLN2 |
| ADK | CACNA1C | EGFL7 | ILK | LIG3 | MIR212 | MIR592 | PPARG | TLX2 |
| ADRB1 | CACNA1D | EIF1 | IMMP1L | LIPA | MIR214 | MIR599 | PPP1R1A | TMEM43 |
| ADRB2 | CACNA1G | EIF4H | IMPG1 | LMNA | MIR216A | MIR615 | PRKAG2 | TMPO |
| AGGF1 | CACNA1H | ELMO3 | ITPR2 | LPL | MIR216B | MIR653 | PSEN1 | TNNC1 |
| AKAP9 | CACNA1S | EML2 | JAG1 | LRAT | MIR217 | MIR758 | PSEN2 | TNNI3 |
| ALMS1 | CACNA2D1 | ENPP1 | JPH2 | LRP6 | MIR22 | MIR761 | PTPN11 | TNNT2 |
| AMPH | CACNA2D2 | ESR1 | JUP | LYRM5 | MIR25 | MIR873 | RAB14 | TPM1 |
| ANK1 | CACNB2 | ESR2 | KCNA4 | LYZ | MIR300 | MIR874 | RABEP2 | TPSG1 |
| ANK2 | CALCR | EVI2A | KCNA5 | MADD | MIR30A | MIR96 | RAF1 | TRPC1 |
| ANKRD1 | CALR | EVI2B | KCNAB1 | MAP2K1 | MIR30B | MKRN2 | RASA1 | TRPC3 |
| AP3M1 | CAMK2A | EYA4 | KCNAB2 | MAP2K2 | MIR31 | MLL | RBM20 | TRPC4 |
| APOA1 | CAMK2B | FAM117A | KCND2 | MCM7 | MIR32 | MRPL23 | RNF130 | TRPC6 |
| APOA2 | CAMK2D | FAS | KCND3 | MED1 | MIR326 | MUTED | RRP8 | TRPC7 |
| APOB | CAMK2G | FBN1 | KCNE1 | MED13L | MIR328 | MYBPC3 | RTL1 | TRPM1 |
| APOC2 | CASQ2 | FGA | KCNE2 | MEST | MIR330 | MYH11 | RYR2 | TRPM3 |
| APOC3 | CAV1 | FKBP2 | KCNH2 | MEX3A | MIR335 | MYH6 | SC5DL | TTN |
| APOC4 | CAV2 | FKTN | KCNH6 | MIB1 | MIR338 | MYH7 | SCN1B | TTR |
| APOE | CAV3 | FLT4 | KCNIP1 | MIR100 | MIR340 | MYH7B | SCN4B | TXNDC5 |
| APP | CCDC73 | FOXC2 | KCNIP2 | MIR107 | MIR346 | MYL2 | SCN5A | VAMP8 |
| ARID3B | CCNH | FOXN3 | KCNJ11 | MIR126 | MIR34A | MYL3 | SETD6 | VANGL1 |
| ARRB1 | CDK14 | FUBP1 | KCNJ12 | MIR132 | MIR34C | MYLK | SFXN5 | VCAN |
| ASTN1 | CETP | GAA | KCNJ2 | MIR134 | MIR367 | MYLK2 | SGCD | VCL |
| ATG10 | CFTR | GALE | KCNJ3 | MIR136 | MIR375 | MYO6 | SHOC2 | VKORC1 |
| ATOH8 | CHCHD4 | GATA4 | KCNJ5 | MIR137 | MIR376B | MYOZ2 | SLC2A10 | VPS13B |
| ATP1A1 | CHD5 | GATA6 | KCNK1 | MIR140 | MIR376C | MYPN | SLC6A2 | VTI1A |
| ATP1A2 | CHRM2 | GBA | KCNK10 | MIR141 | MIR377 | NCRNA00081 | SLC9A1 | WIT1 |
| ATP1A3 | CLCA4 | GGCX | KCNK13 | MIR143 | MIR382 | NDE1 | SNAPC5 | WNK1 |
| ATP1A4 | CLCN3 | GINS3 | KCNK16 | MIR144 | MIR410 | NEXN | SNORA46 | WT1 |
| ATP1B1 | CNOT1 | GJA1 | KCNK17 | MIR145 | MIR411 | NF1 | SNORA50 | WWP2 |
| ATP1B2 | COL27A1 | GJA5 | KCNK3 | MIR149 | MIR425 | NISCH | SNTA1 | ZFPM2 |
| ATP1B3 | COL3A1 | GNB1L | KCNK5 | MIR150 | MIR429 | NKX2-5 | SOS1 | ZNF221 |
| ATP2A1 | COL4A1 | GPC1 | KCNK6 | MIR152 | MIR431 | NME7 | SSH2 | ZNF225 |
| ATP2A2 | COL4A2 | GPD1L | KCNK9 | MIR154 | MIR433 | NOS1AP | STAMBPL1 | ZNF230 |
| ATP2A3 | COPZ2 | GRID1 | KCNMB1 | MIR155 | MIR451 | NOTCH1 | STARD3 | ZNF283 |
| ATP2B1 | CRYAB | GRM8 | KCNN3 | MIR182 | MIR455 | NPHP4 | SUN1 | ZRANB2 |
| ATP2B2 | CSRP3 | GSN | KCNN4 | MIR183 | MIR484 | NPPA | SUN2 | |
| BCL2L14 | CST3 | GTPBP1 | KCNQ1 | MIR185 | MIR485 | NRD1 | SYNE1 | |
| BIN1 | CTF1 | HCN1 | KIAA1797 | MIR186 | MIR486 | OMG | SYNE2 | |
| BMP2 | CUX2 | HCN2 | KLHL3 | MIR190 | MIR487B | OPN4 | TAF10 | |
| BMP4 | DALRD3 | HCN4 | KRAS | MIR195 | MIR488 | OSMR | TARS2 | |
| BMP6 | DBH | HMGCL | KRT14 | MIR197 | MIR489 | PANK1 | TBX1 | |
| BMPR2 | DES | HMGCS2 | KRT16 | MIR200A | MIR490 | PCSK9 | TBX20 | |
| BRAF | DHCR24 | HOPX | KRT17 | MIR202 | MIR491 | PDE1A | TBX5 | |
| Supplementary Table 2: Genes known to be associated with human cardiovascular disease. | | | | | | | | |

### Slide 4
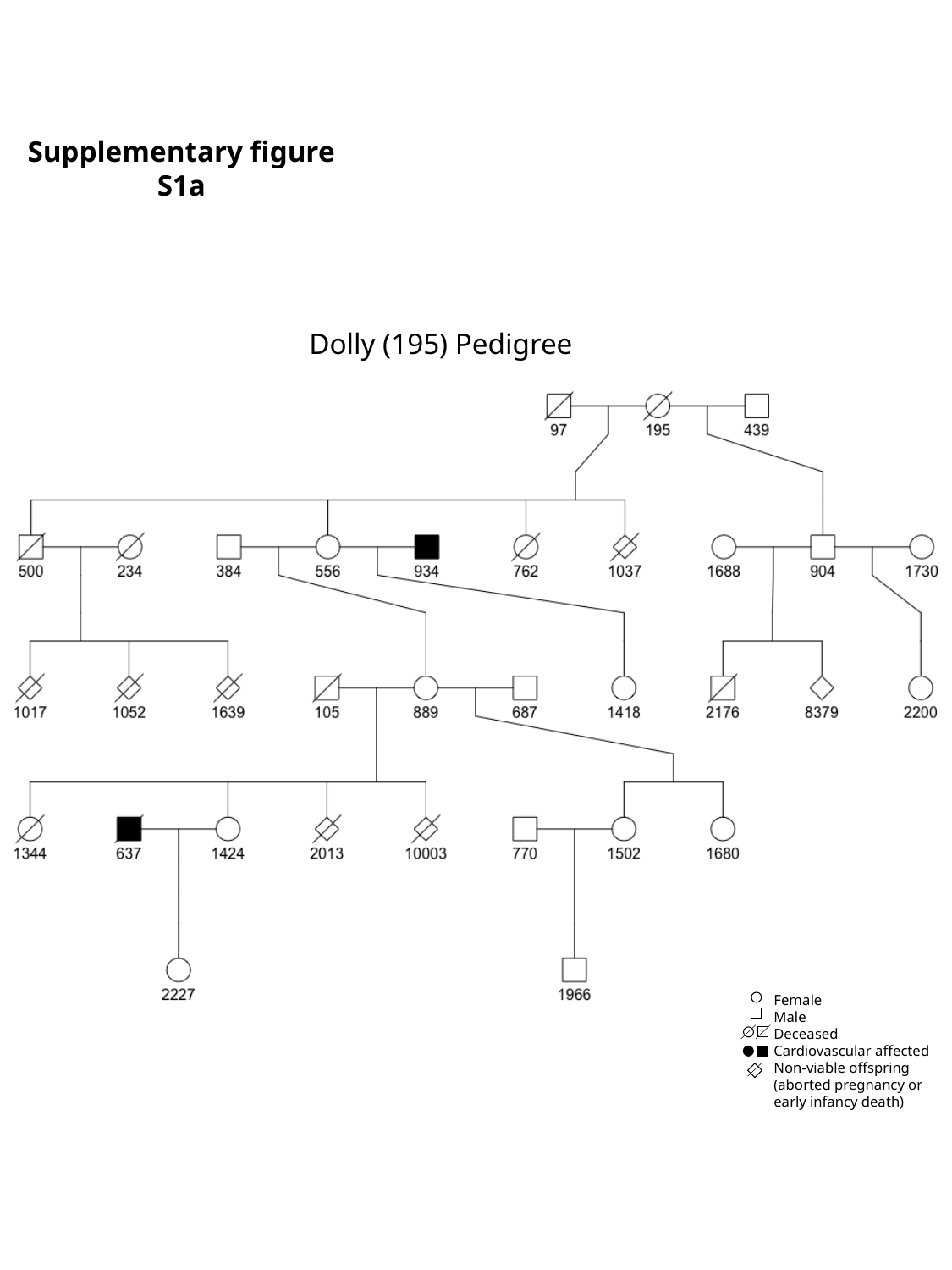

Supplementary figure S1a
Dolly (195) Pedigree
Female
Male
Deceased
Cardiovascular affected
Non-viable offspring (aborted pregnancy or early infancy death)

### Slide 5
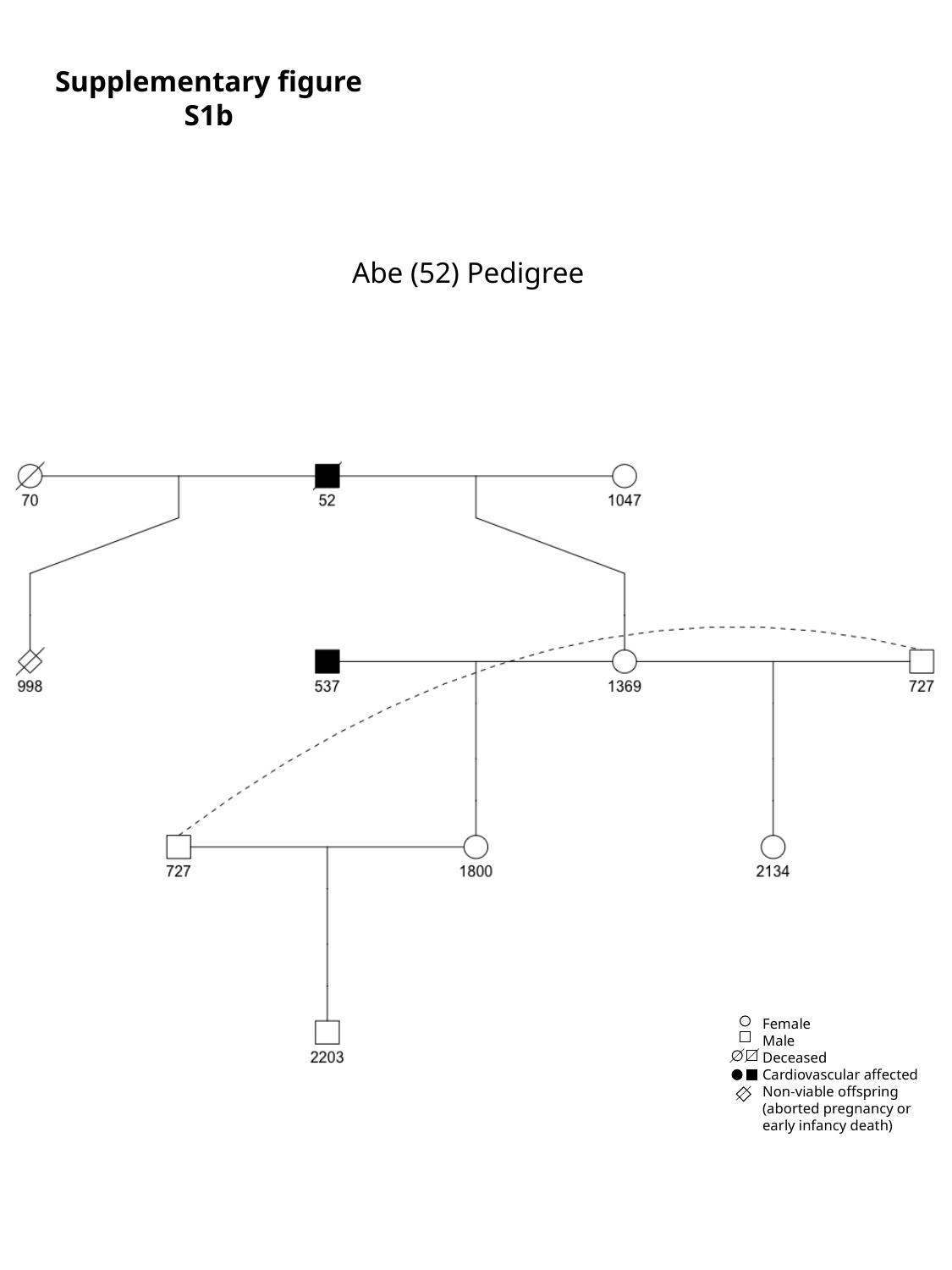

Supplementary figure S1b
Abe (52) Pedigree
Female
Male
Deceased
Cardiovascular affected
Non-viable offspring (aborted pregnancy or early infancy death)

### Slide 6
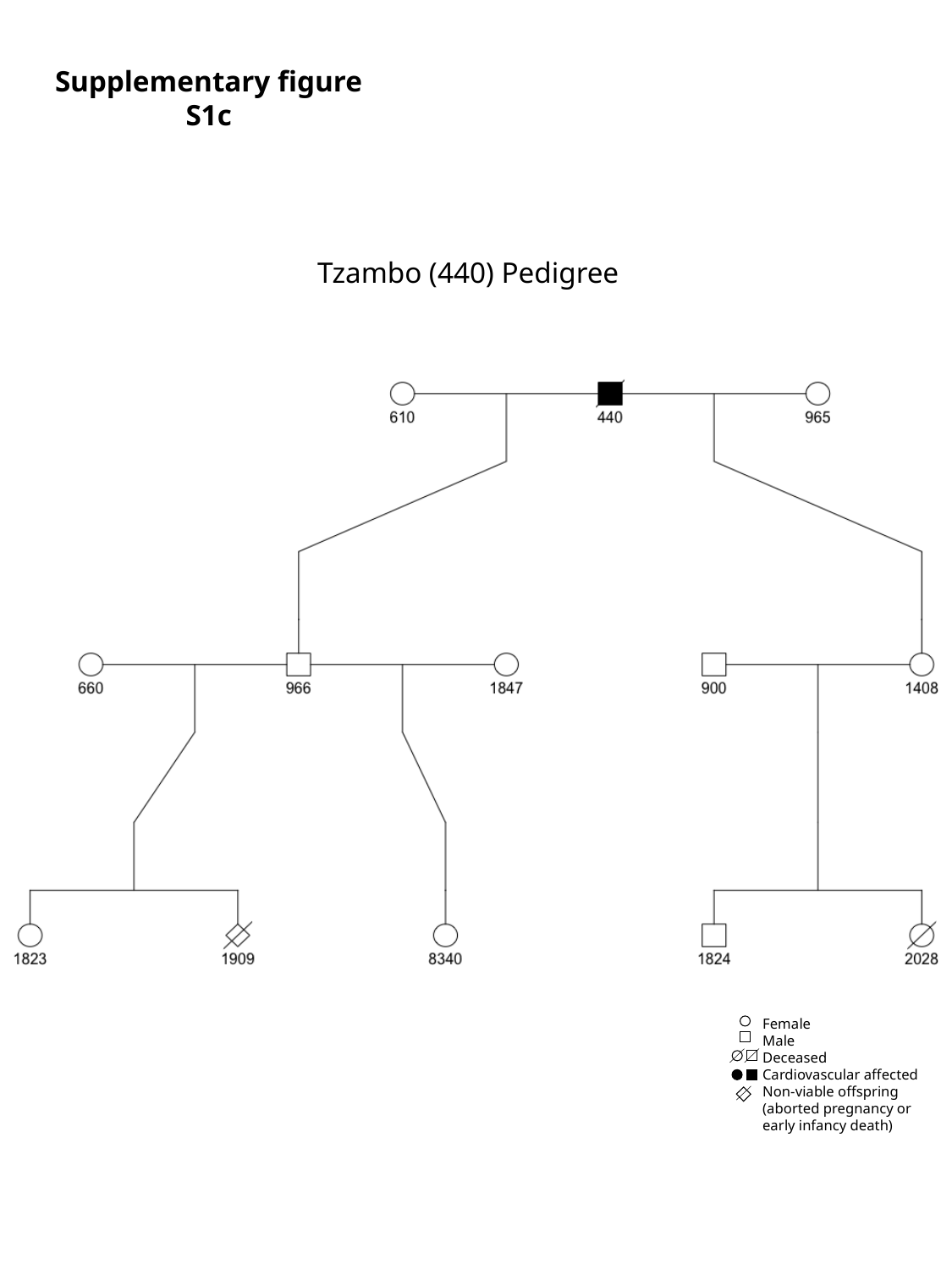

Supplementary figure S1c
Tzambo (440) Pedigree
Female
Male
Deceased
Cardiovascular affected
Non-viable offspring (aborted pregnancy or early infancy death)

### Slide 7
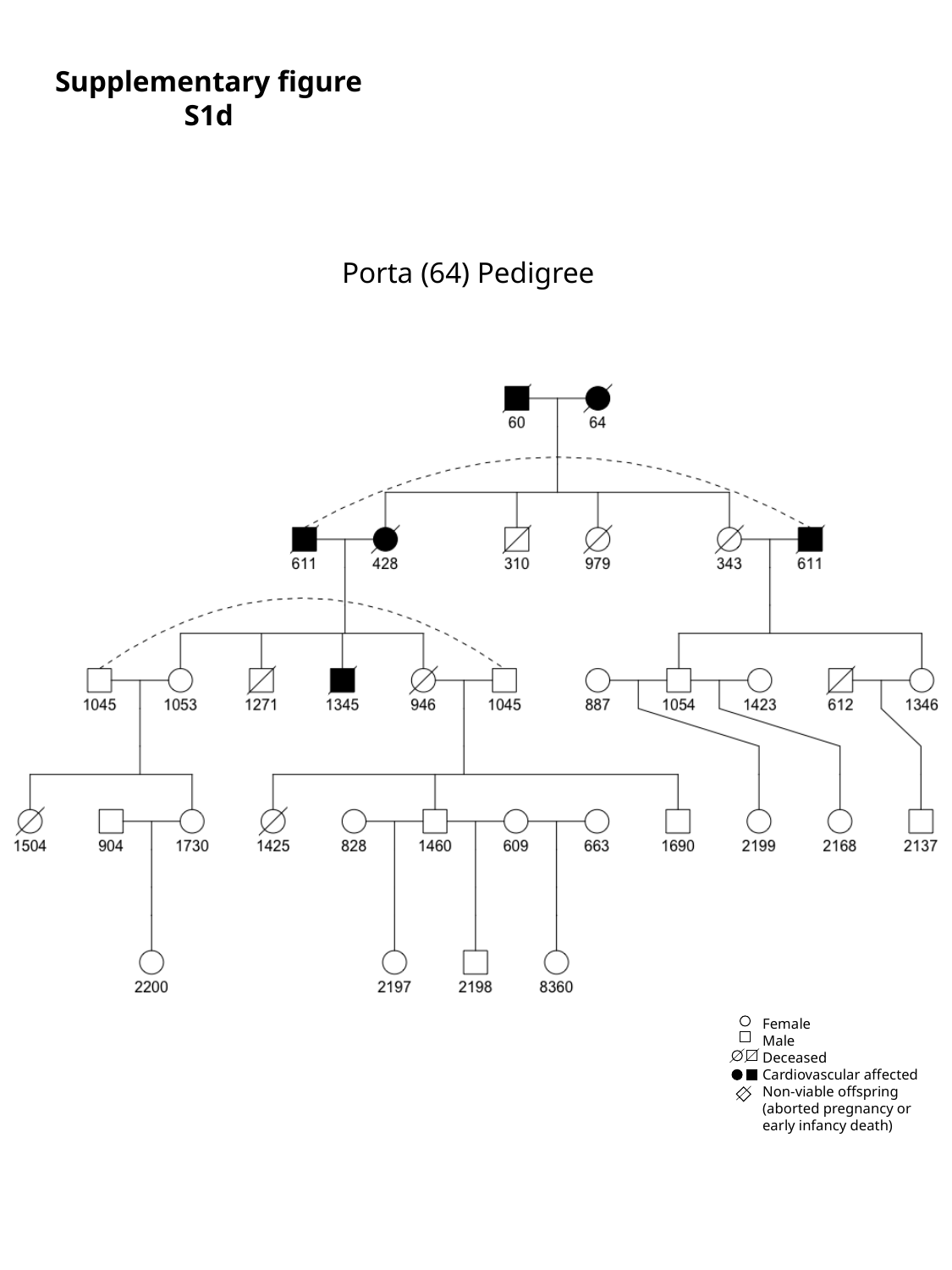

Supplementary figure S1d
Porta (64) Pedigree
Female
Male
Deceased
Cardiovascular affected
Non-viable offspring (aborted pregnancy or early infancy death)

### Slide 8
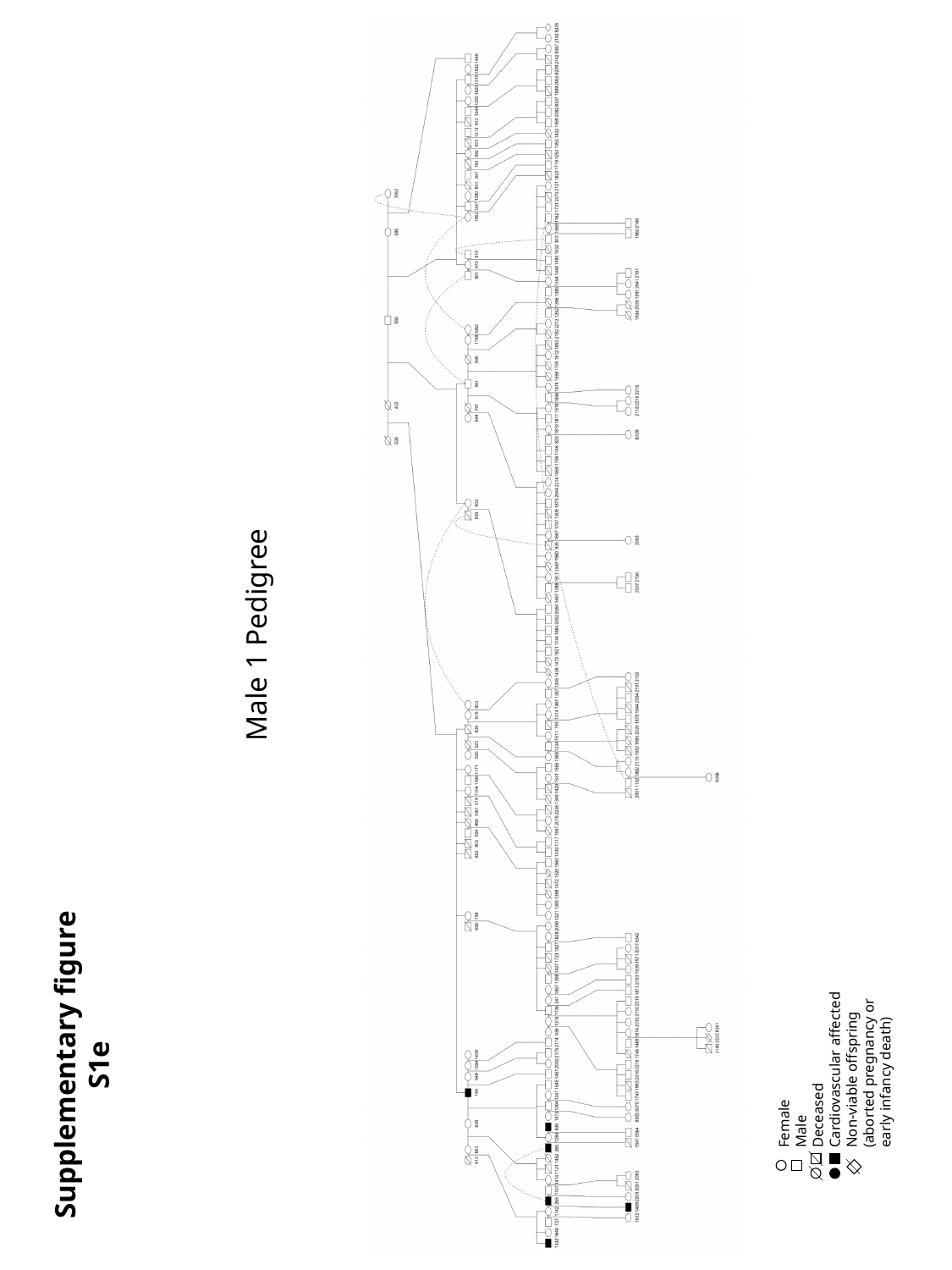

Male 1 Pedigree
Female
Male
Deceased
Cardiovascular affected
Non-viable offspring (aborted pregnancy or early infancy death)
Supplementary figure S1e

### Slide 9
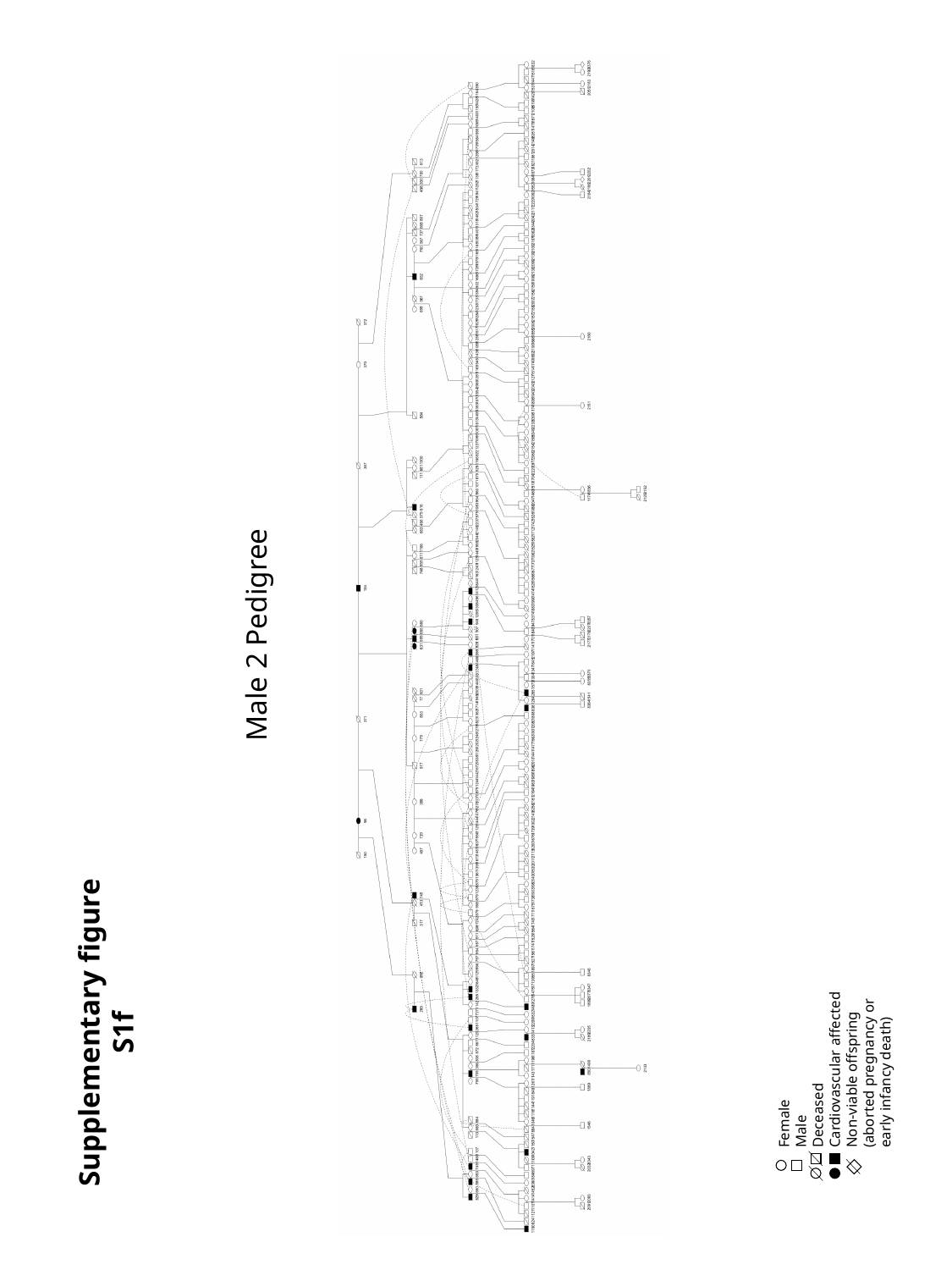

Male 2 Pedigree
Female
Male
Deceased
Cardiovascular affected
Non-viable offspring (aborted pregnancy or early infancy death)
Supplementary figure S1f

### Slide 10
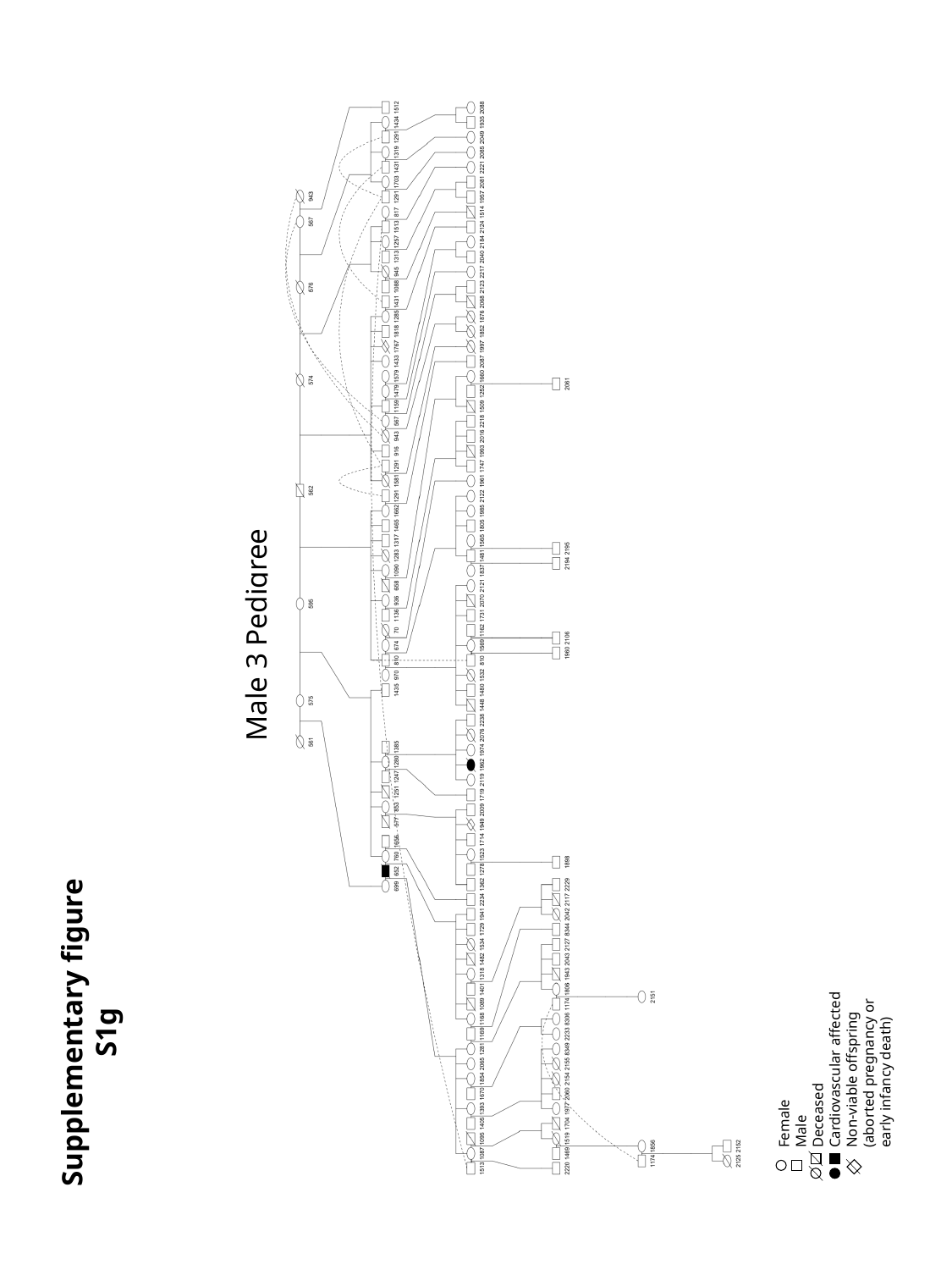

Male 3 Pedigree
Supplementary figure S1g
Female
Male
Deceased
Cardiovascular affected
Non-viable offspring (aborted pregnancy or early infancy death)

### Slide 11
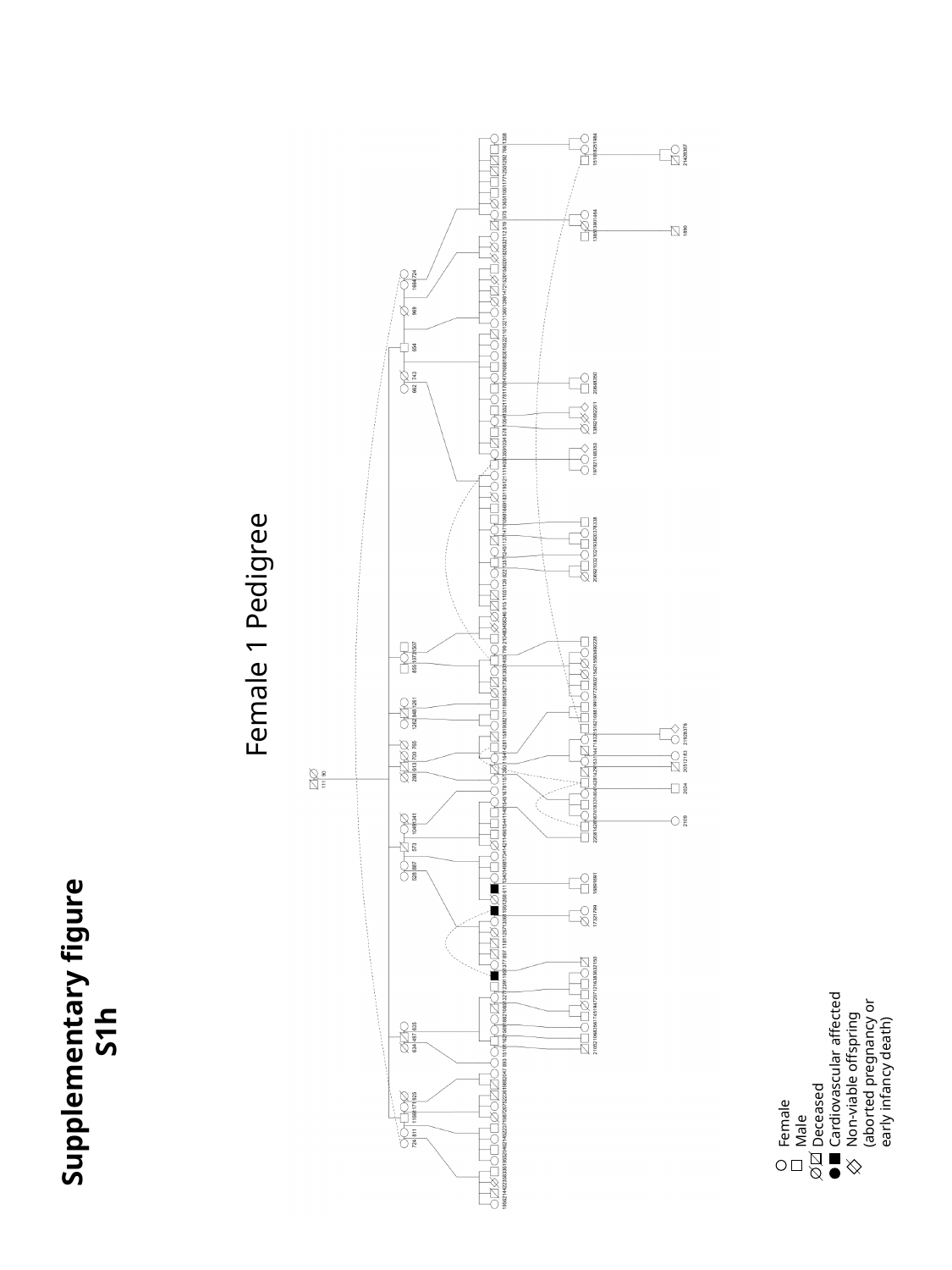

Female 1 Pedigree
Supplementary figure S1h
Female
Male
Deceased
Cardiovascular affected
Non-viable offspring (aborted pregnancy or early infancy death)

### Slide 12
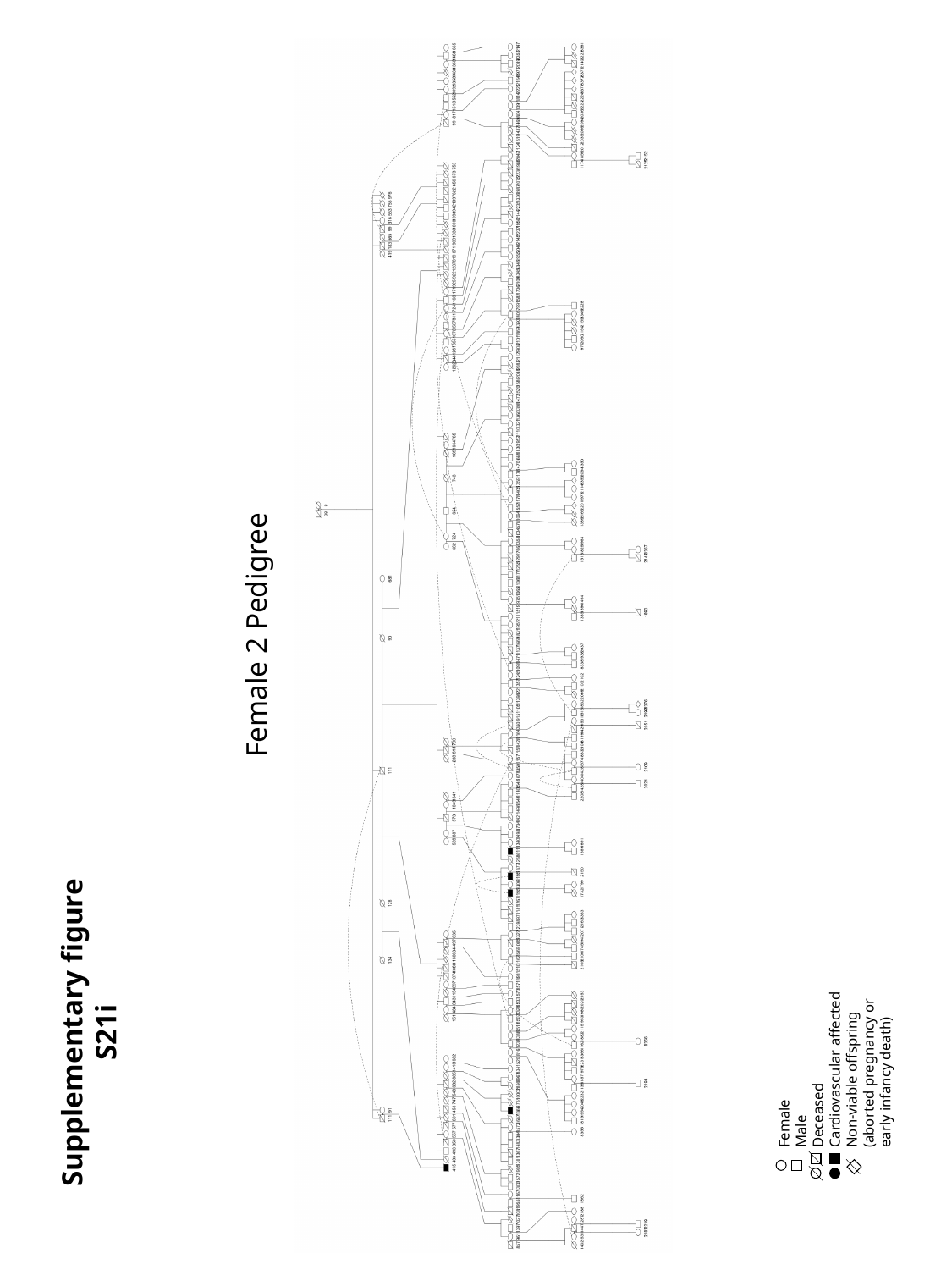

Female 2 Pedigree
Supplementary figure S21i
Female
Male
Deceased
Cardiovascular affected
Non-viable offspring (aborted pregnancy or early infancy death)

### Slide 13
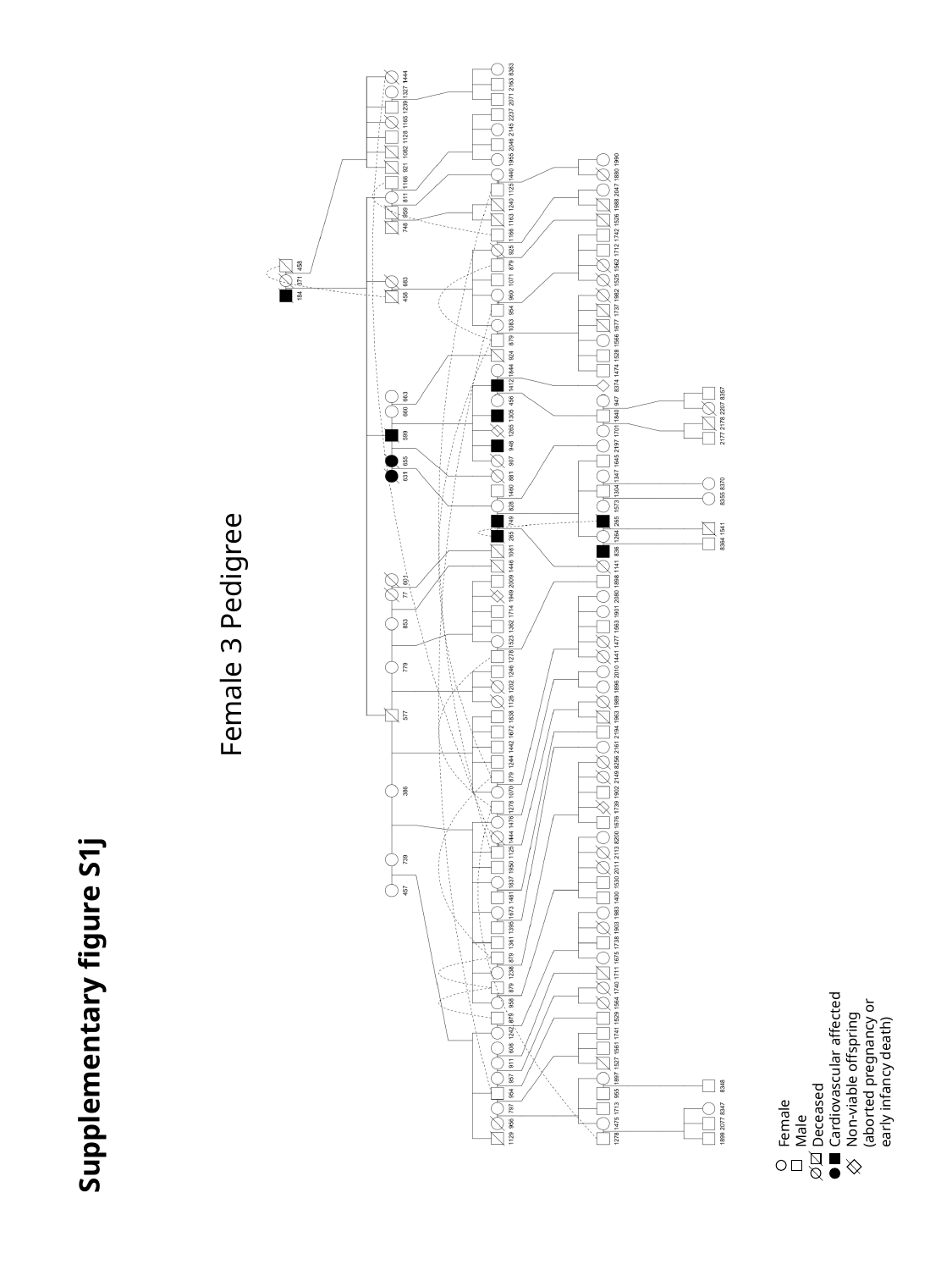

Female 3 Pedigree
Supplementary figure S1j
Female
Male
Deceased
Cardiovascular affected
Non-viable offspring (aborted pregnancy or early infancy death)
