## Supplementary Text for "Leveraging whole genome sequencing to promote genetic diversity and population health in zoo-housed western lowland gorillas"

### Use of Kinship Values to promote genetic diversity in zoo populations

Genetically diverse populations are more likely to adapt to environmental changes and resist disease.^39,40^ However, breeding within a closed population--defined as a population that has no potential for external genetic contribution--will result in a decrease of genetic diversity as a result of genetic drift and inbreeding.^41^ In order to minimize the loss of genetic diversity the SSP and EEP, breeding priority is given to those individuals who share the least kinship with each other and have the least individual mean kinship to the general population.^42^ Of these two measurements the kinship coefficient (*k_i_*) is defined as the probability that alleles randomly selected from homologous loci in two individuals (*P_a_ , P_b_*) are identical by descent,

and the individual mean kinship (*mk_i_*), is used to identify the mean relatedness an individual shares with the entire population (*K_ab_*=kinship shared between individual a and b, *N*=total number of individuals in the population).
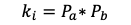

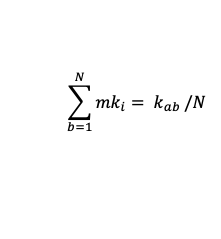

Determination of kinship coefficients from the western lowland gorilla population have family structure (pedigree) information held in the international studbook for the western lowland gorilla (held by the AZA, EZA and dewarwildlife.org). These record-keeping documents contain the pedigrees of all zoologic housed western lowland gorillas. However, lineage of zoologic housed gorillas can only be traced back to the founding individuals defined as wild born individuals who were introduced to captivity. While the region and year of capture of the wild born gorillas was recorded, no detailed information regarding family structure is known. Identification of family structure based on location and date of capture is expected to be inaccurate, and therefore, wild-born individuals are assumed to be unrelated within the studbook. A potential consequence of this assumption includes mating of related gorillas resulting in consanguinity of offspring and a reduced genetic diversity within the *ex situ* population.

### Health Information of the Gorillas Used for Segregation Analysis

Dolly, a female wild-born western lowland gorilla was housed at the San Diego Zoo (1963 - 1988) where she died at the estimated age of 26 years. Her cause of death was Toxemia/septicemia as a result of a small intestinal volvulus due to herniation through a mesenteric rent. Necropsy Results showed no indication of myocardial fibrosis. Dolly had four viable offspring during her lifetime and a complete pedigree is available (Supplementary table S2a)

Abe, a male wild-born western lowland gorilla was housed in Colorado Springs (1956-1971), Brownsville (1971-1972), Colorado Springs (1972-1979), San Diego Zoo (1979-1981), Houston (1979-1981), Topeka (1981-1984), Omaha (1984-1992), Chicago Brookfield (1992-1995) where he died at the approximate age of 39 years. Abe’s cause of death was reported to be chronic pneumonia and suppurative periodontitis. Necropsy results indicate interstitial myocardial fibrosis. Abe had one viable offspring during his lifetime and a complete pedigree is available (Supplementary Table S2b)

Tzambo, a male wild-born western lowland gorilla was housed in Weybridge (1972-1984), San Diego (1984-1985), and Los Angeles (1985-1999) where he died at the approximate age of 26 years. His cause of death was reported to be an aortic aneurysm, with evidence of dilated cardiomyopathy. Necropsy results suggest interstitial myocardial fibrosis. Tzambo had 2 viable offspring during his lifetime and a complete pedigree is available (Supplementary Table S2c).

Porta, a female wild-born western lowland gorilla was housed in Toledo (1957-1993) where she died at the approximate age of 39 years. Porta’s cause of death was euthanasia as a result of end-stage heart failure. Necropsy results suggest interstitial myocardial fibrosis. Porta had four viable offspring during her lifetime and a complete pedigree is available (Supplementary Table S2
